# Demographic effects of aggregation in the presence of a component Allee effect

**DOI:** 10.1101/2023.05.12.540532

**Authors:** Daniel C.P. Jorge, Ricardo Martinez-Garcia

## Abstract

Intraspecific interactions are key drivers of population dynamics because they establish relations between individual fitness and population density. The component Allee effect is defined as a positive correlation between any fitness component of a focal organism and population density, and it can lead to positive density dependence in the population per capita growth rate. The spatial population structure is key to determining whether and to which extent a component Allee effect will manifest at the demographic level because it determines how individuals interact with one another. However, existing spatial models to study the Allee effect impose a fixed spatial structure, which limits our understanding of how a component Allee effect and the spatial dynamics jointly determine the existence of demographic Allee effects. To fill this gap, we introduce a spatially-explicit theoretical framework where spatial structure and population dynamics are emergent properties of the individual-level demographic and movement rates. Depending on the intensity of the individual-level processes the population exhibits a variety of spatial patterns, including evenly spaced aggregates of organisms, that determine the demographic-level by-products of an existing individual-level component Allee effect. We find that aggregation increases population abundance and allows populations to survive in harsher environments and at lower global population densities when compared with uniformly distributed organisms. Moreover, aggregation can prevent the component Allee effect from manifesting at the population level or restrict it to the level of each independent group. These results provide a mechanistic understanding of how component Allee effects might operate for different spatial population structures and show at the population level. Because populations subjected to demographic Allee effects exhibit highly nonlinear dynamics, especially at low abundances, our results contribute to better understanding population dynamics in the presence of Allee effects and can potentially inform population management strategies.

## 1 Introduction

Intraspecific interactions are critical to understanding population ecology because they define how demographic rates depend on population density and ultimately drive population dynamics. The Allee effect is characterized by a positive correlation between population size or density and any individual fitness component (Courchamp et al., 2008; Levitan, 2005; Stephens et al., 1999). Because of this positive density dependence, populations subjected to Allee effects might have thresholds for population survival that manifest in sudden extinctions, existence of alternative stable states, and hysteresis (Courchamp et al., 2008; Lande, 1987; Oro, 2020a; Sun, 2016). These highly nonlinear features make populations exhibiting Allee effects hard to manage without a mechanistic understanding of how the individual-level processes and interactions that underlie the Allee effect are responsible for the trends and patterns observed in population dynamics.

Allee effects are studied mainly at two levels: the component and the demographic Allee effect (Stephens et al., 1999). The component Allee effect is a positive association between population density and one (or many) components of individual fitness, such as offspring survival, mating success, or fecundity (Courchamp et al., 2008; Drake and Kramer, 2011; Orr, 2009) (Fig. 1a). Component Allee effects rely on several mechanisms. In some fish, rotifer, and mammals such as marmots, the presence of conspecifics changes the environmental conditions locally, improving habitat quality and individual fitness (Allee and Bowen, 1932; Allee and Rosenthal, 1949; Ghazoul, 2005; Stephens et al., 2002). Especially in group-living organisms, cooperative behaviors such as group vigilance, nursing, resource sharing, and social foraging also make individuals more competent in the presence of conspecifics (Angulo et al., 2018, 2013; Dechmann et al., 2010; Luque et al., 2013; Nowak and Lee, 2011; Snaith and Chapman, 2008). Allee effects are also frequent in sexually reproducing species. In motile organisms, females are more likely to find mates at larger population sizes (Dennis, 1989; Garrett and Bowden, 2002; Liermann and Hilborn, 2001; Tcheslavskaia et al., 2002). In sessile organisms, such as pollinators or broadcast spawners, fecundation is more likely at high population densities (Ashman et al., 2004; Guy et al., 2019; Lundquist and Botsford, 2011; Luzuriaga et al., 2006; Wagenius, 2006). On the other hand, the demographic Allee effect is a population-level emergent property due to the existence of one or more component Allee effects, and it manifests as a positive correlation between the net per-capita growth rate and the population size. This positive density-dependence is easier to identify at low population densities because competition hinders its effect in more crowded scenarios (Courchamp et al., 2008). Demographic Allee effects are strong if the population cannot survive below a specific threshold size (Allee threshold) or weak if positive density-dependence is not intense enough to establish such a survival threshold (Courchamp et al., 2008; Drake and Kramer, 2011).

**Figure 1:**
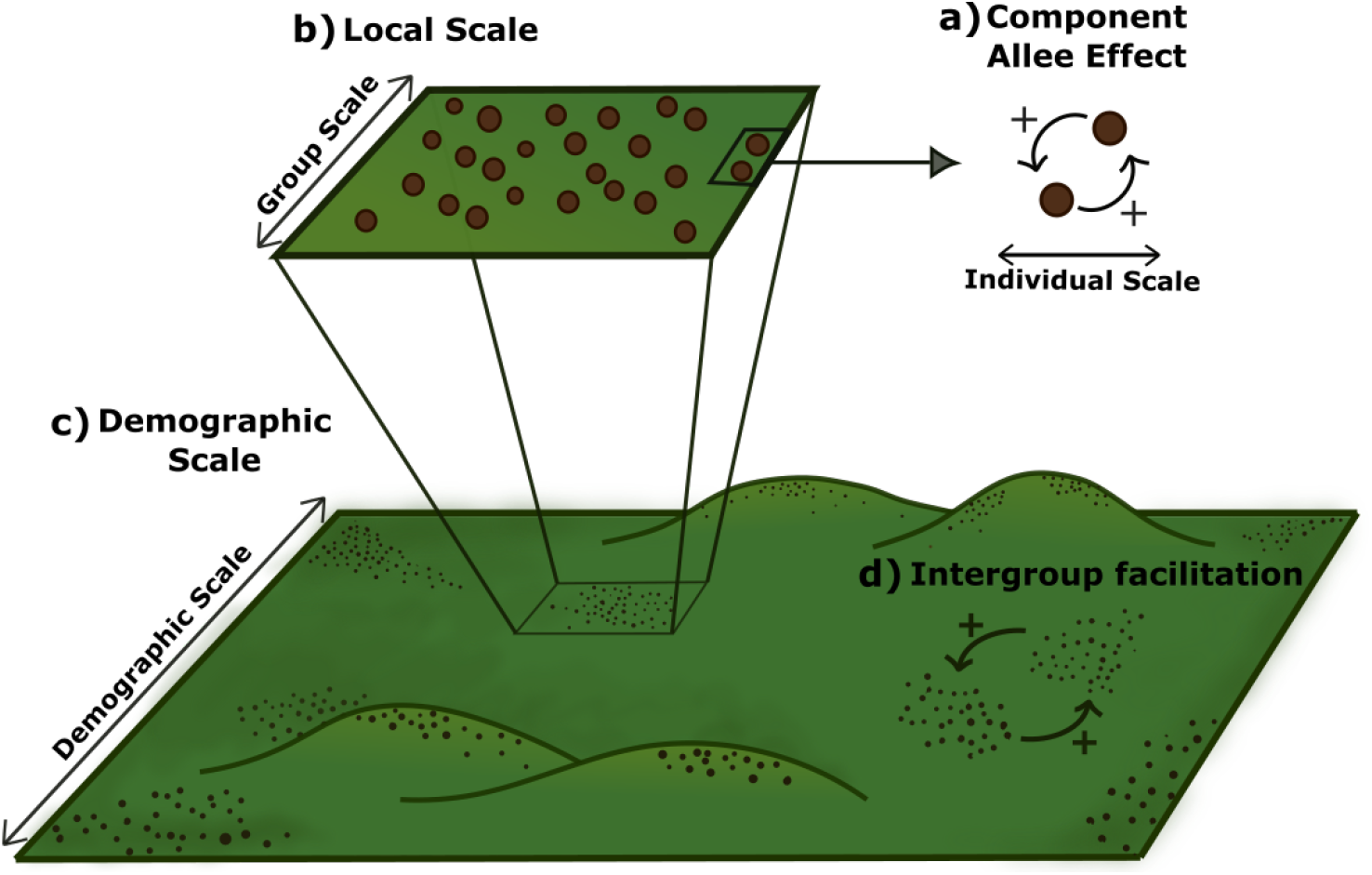
Allee effect across spatial scales. The component Allee effect (a) is a result of interactions between individuals that manifests at a (b) local scale around a focal organism. At the demographic scale (c), individuals are spatially scattered, possibly forming aggregates. In the presence of aggregates, the population has a fourth characteristic scale, defining inter-group facilitation (d)

The fitness of a focal individual in the presence of a component Allee effect is a nonlinear function of the local density of conspecifics around it (Fig. 1b). Moreover, because Allee effects have a more substantial impact at low population densities and often require the direct interaction between at least two organisms, the spatial population structure is key to determining whether and to which extent a component Allee effect will manifest at the demographic level (Kanarek et al., 2013; Kramer et al., 2009; Surendran et al., 2020) (Fig. 1c). Back to Allee’s seminal experiments, several studies have investigated the impact of the spatial population structure, and more specifically of aggregation, on Allee effects (Allee, 1938). For instance, some plant populations produce more and heavier seeds if distributed in clumps (Luzuriaga et al., 2006; Wagenius, 2006). Plant aggregates can also facilitate nearby individuals because they attract pollinators to them, which extends the facilitation range beyond the scale of a single cluster of plants (Fig. 1d) (Le Cadre et al., 2008), and ameliorate physical stresses (Silliman et al., 2015). Broadcast spawners subjected to a strong Allee effect, such as the red sea urchin *Strongylocentrotus franciscanus*, can survive at low abundances by aggregating (Guy et al., 2019; Lundquist and Botsford, 2011). Finally, several social species form spatially segregated groups, which could contribute to population persistence in harsh environmental conditions (Angulo et al., 2018; Lerch et al., 2018; Woodroffe et al., 2020). Aggregation and group living are thus ubiquitous features of populations subjected to Allee effects, and they strongly influence the emergent population dynamics. To explain how these spatial features impact populations subjected to component Allee effects, recent studies have introduced the group-level Allee effect, defined as any positive association between the organism’s fitness and group size (Lerch et al., 2018). However, a theoretical framework describing how group-level Allee effects emerge from component Allee effects and the individual-level processes responsible for aggregation and group formation is lacking.

Over the last decades, theoretical studies have been key to develop much of our current understanding of Allee effects (Asmussen, 1979; Cushing, 1988; Hsu and Fredrickson, 1975; Kostitzin, 1940; Lande, 1987; Sun, 2016; Tammes et al., 1964; Volterra, 1938). Several models, either deterministic or stochastic, consider well-mixed populations and disregard spatial degrees of freedom (Dennis, 1981, 2002; Méndez et al., 2019). The effect of space has been investigated mainly using metapopulation approaches in which each node represents a group or cluster of individuals and links represent any inter-group interaction (Padrón and Trevisan, 2000; Rijnsdorp and Vingerhoed, 2001). These frameworks already incorporate group-level Allee effects because they restrict fitness benefits due to intraspecific interactions to each metapopulation and have helped explain why component Allee effects rarely many manifest at the demographic level in group-living species (Courchamp et al., 2008; Rijnsdorp and Vingerhoed, 2001). However, metapopulation models impose the existence of groups in the stationary state and do not describe the group-forming dynamics. Alternative approaches, based on individual-based models (IBMs) or partial differential equations (PDEs), incorporate space explicitly and can describe the group-forming dynamics (Keitt et al., 2001; Maciel and Lutscher, 2015; Surendran et al., 2020; Wang et al., 2019). Therefore, these approaches can explain how different spatial patterns of population density impact the outcome of ecological dynamics, such as species invasions, in the presence of Allee effects (Keitt et al., 2001; Maciel and Lutscher, 2015) or Allee-effect features, such as the Allee threshold (Surendran et al., 2020).

In this work, we develop a theoretical framework to investigate Allee effects across different levels of spatial organization within a population. We present this formalism starting from a stochastic and spatially explicit individual-based description of a population with density-dependent reproduction mimicking a component Allee effect. This description is the most fundamental level at which we can describe a population, allowing us to explicitly model the relationship between the mechanism responsible for the component Allee effect and individual birth and death rates. From this individual-level description, we derive the corresponding deterministic equation for the dynamics of the population density. This approximation allows us to investigate in which conditions individuals aggregate due to individual-level interactions and to study the population-level consequences of the component Allee effect depending on the spatial population structure. Finally, we identify the cases in which we can describe the long-term spatial distribution of individuals in terms of a metapopulation model, and use this approach to investigate the emergence of group-level Allee effects. Our results recapitulate several observations on the interplay between spatial structure, group, and demographic Allee effects, providing a unifying theoretical framework to investigate the interplay between component Allee effects and spatial dynamics.

## 2 Methods

### 2.1 A spatially explicit individual-based model with component Allee effect

At the most fundamental level, we describe the spatio-temporal population dynamics using an IBM in which we can incorporate any ecological interaction, such as competition, predation, or cooperation, movement, and birth-death dynamics tracking single individuals. We consider a population with density-independent birth, death, and movement and also account for density-dependent birth and death processes. Specifically, individuals interact via binary reproductive facilitation and ternary competition. Reproductive facilitation is common even in species with asexual reproduction when individuals need the presence of conspecifics to reach the physiological condition to reproduce (Courchamp et al., 2008). Some examples of species exhibiting asexual reproduction and reproductive facilitation are self-fertile snails, and parthenogenetic female lizards (Crews et al., 1986; Thomas and Benjamin, 1974). Competition, on the other hand, reduces individual fitness at very high population densities and is necessary to avoid unbounded population growth. The combination of binary reproductive facilitation and ternary competition results in a hump-shaped relationship between per capita reproduction rate and local density of individuals similar to that reported by Allee in his experiments with laboratory populations of the flour beetle (Allee, 1938; Allee et al., 1949).

We can summarize the previous processes and interactions in the following set of demographic reactions

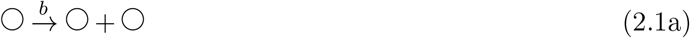

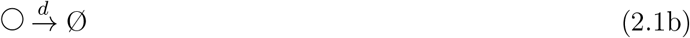

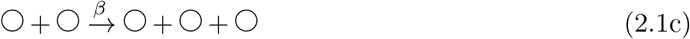

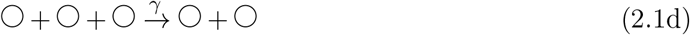

(2.1a) and (2.1b) represent density-independent birth and death. (2.1c) represents a binary cooperative interaction in which two individuals interact at rate *β* and produce a third individual. The last reaction, (2.1d), describes ternary competition. This set of processes is one of the mathematically simplest ways of modeling a component Allee effect at the individual level (Méndez et al., 2019). However, one can think of many other density-dependent processes that might result in a component Allee effect, such as reduced death, sexual reproduction, or collective predation, among others (Drake and Kramer, 2011; Oro, 2020b). Any of these alternative processes can be incorporated into our modeling approach by simply modifying the set of reactions (2.1).

To introduce spatial dynamics, we consider that individuals are located in the sites of a onedimensional regular lattice with periodic boundary conditions, but it is straightforward to extrapolate the derivation to more realistic two-dimensional landscapes. We label each lattice node with an integer index *i* ∈ [0, *N*], and denote the spatial coordinate with *x* ∈ [0, *L*]. The distance between two adjacent lattice nodes is *δx* such that the spatial coordinate of the *i*-th node is *x*_*i*_ = *i δx*. Individuals move on the lattice performing a nearest-neighbor random walk, and the density-dependent interactions in (2.1c)-(2.1d) only occur if individuals are within an interaction-specific range.

We can express individual random movement using the reaction notation of (2.1) as

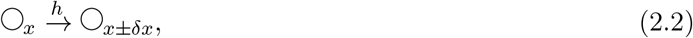

where *h* is the jump transition rate and *δx* is the displacement length. These choices result in a diffusive movement with diffusion coefficient *D* = *h δx*^2^. To account for the spatial extent of the interactions between individuals, we modify the demographic rates in the reactions (2.1c) and (2.1d). We consider that two individuals facilitate one another if they are closer than the facilitation range *R*_*f*_. As a result, a focal individual at location *x* reproduces with rate *β/*2*R*_*f*_. In terms of reactions, this process can be written as:

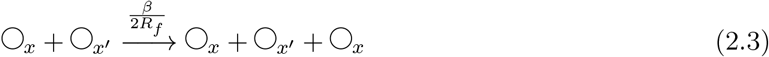

provided that |*x−x*′| ≤ *R*_*f*_. For negative interactions, we consider that a focal individual at location *x* can die due to competition by forming triplets with two neighbors at locations *x*′ and *x*″. This process occurs with rate 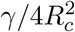 provided that the distance between the focal individual and each of these two neighbors is shorter than or equal to the competition range |*x−x*′| ≤ *R*_*c*_ and |*x−x*″| ≤ *R*_*c*_. In terms of reactions, we can write this process as

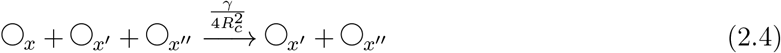

Finally, notice that both for the facilitation and the competition terms, we are assuming that the non-local reaction rates do not depend on the distance between individuals as long as the pairwise distances between individuals in a pair or triplet are shorter than the interaction range. We are therefore modeling the interaction kernel with a top-hat function. The factors dividing the rates *β* and *γ* are normalizing factors of the top-hat kernel. This normalization makes birth/death rates depend on population density rather than on population size.

### 2.2 Derivation of population-level approximations

We use the Doi-Peliti formalism to derive a deterministic approximation of the spatial stochastic dynamics introduced in Section 2.1. This deterministic approximation neglects demographic fluctuations and maps the set of discrete reactions to a deterministic partial differential equation that describes the dynamics of a population density field *ρ*(*x, t*) in continuous space and time (Doi, 1976; Hernández-García and López, 2004; Peliti, 1985; Täuber, 2007). Hence, this approximation fails to describe noise-driven consequences of the Allee effect that might be ecologically relevant at low population sizes, such as extinctions caused by demographic noise (Méndez et al., 2019). It, however, allows us to apply tools from spatially-extended dynamical systems and obtain analytical insights of the underlying stochastic dynamics. More specifically, we can investigate in which conditions individuals form aggregates, resulting in a regular spatial pattern of population density (Cross and Hohenberg, 1993). Following the steps detailed in the Supplementary Material section S1, the stochastic dynamics defined in Section 2.1 leads to the following partial differential equation for *ρ*(*x, t*)

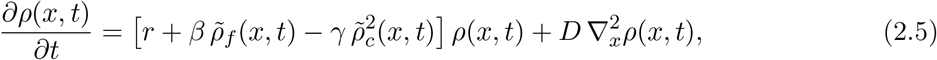

where

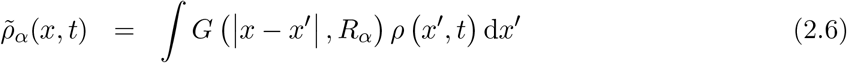

with *α* = {*f, c*} for facilitation and competition, respectively. *G*(|*x − x*′| ; *R*_*α*_) is the normalized interaction kernel for each of the intraspecific interactions

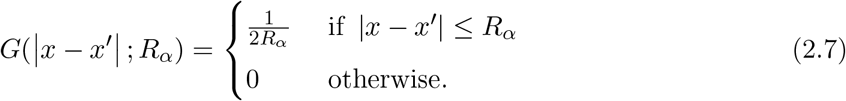

When the population density is uniform, the nonlocal model of Eq. (2.5) is mathematically equivalent to the cubic model used in the literature as the paradigmatic example of a population-level model with demographic Allee effect (Kot, 2001; Méndez et al., 2019; Oro, 2020a). This cubic model has two stable stationary solutions and one unstable. One of the stable stationary solutions is the extinction state. The second stable stationary state, *ρ*_+_, and the unstable one, *ρ*_*−*_, are the roots of the quadratic equation *r* + *β ρ − γ ρ*^2^ = 0,

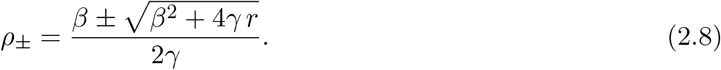

Finally, because *ρ*(*x, t*) is a population density, we must integrate it over the system size to obtain the total population size,

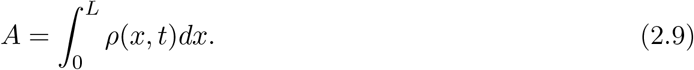

## 3 Results

### 3.1 Group formation

We first perform numerical simulations of the stochastic dynamics represented by the set of reactions in (2.1)-(2.4) using the Gillespie algorithm (Gillespie, 1977). For high diffusion, i.e. high values of *h*, the population reaches a steady state with a uniform spatial distribution of organisms (Fig. 2a, b). As diffusion decreases, however, individuals start to aggregate and the population develops a spatial pattern characterized by isolated clumps of organisms interspersed with unpopulated regions (Fig. 2c-f). Moreover, the total population size increases in the stationary state due to aggregation (Fig. 2g), indicating that grouping improves the environmental conditions and increases the system carrying capacity. The same type of spatial structure and population dynamics are observed in two dimensions (Fig. S1).

**Figure 2:**
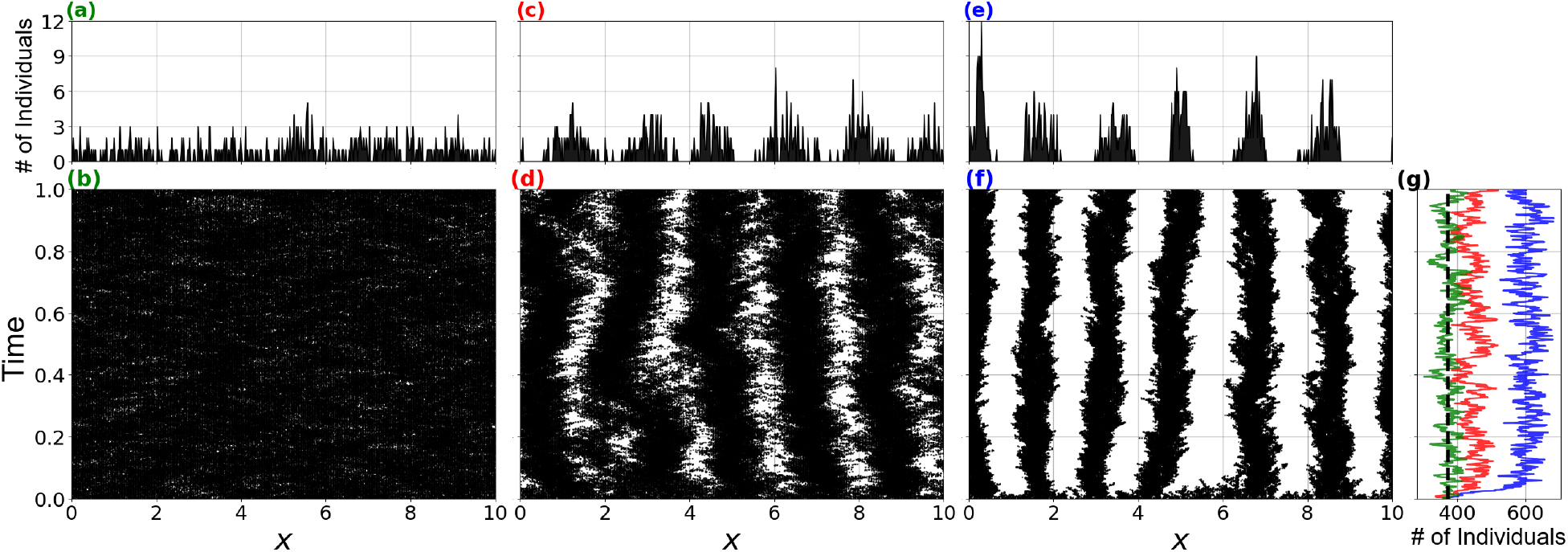
Emergence of spatial patterns for different diffusion regimes. Spatial distribution of individuals resulting from the individual-based stochastic model for (a-b, *D* = 0.08), intermediate (c-d, *D* = 1.2), and high diffusion (e-f, *D* = 8). Top panels (a, c, e) show the number of individuals at each lattice node at the end of a single simulation run. Bottom panels (b, d, f) show the temporal dynamics of the spatial distribution of individuals. The leftmost panel (g) shows the dynamics of population size at high (green), intermediate (red), and low (blue) diffusion together with the prediction from the non-spatial model (black-dashed line), *A* = *ρ*_+_*L*. Bottom panels (b, d, f, g) share the same time scale in the vertical axis and top panels share the same *x* axis as their bottom counterparts. Other parameter values for all panels: *b* = 30, *d* = 40, *β* = 4, *γ* = 0.1, *L* = 10, *R*_*f*_ = 0.75 and *R*_*c*_ = 1, *δx* = 0.02; uniform initial condition. See Supplementary Material section S6 for details on the algorithm.

Next, we compare these simulation outcomes with the results of integrating the deterministic approximation in Eq. (2.5). Our results return a very good quantitative agreement between the stochastic individual-level dynamics and the deterministic equation for population density (Fig. 3), which allows us to use the latter to investigate in which conditions aggregates form and their population-level consequences.

**Figure 3:**
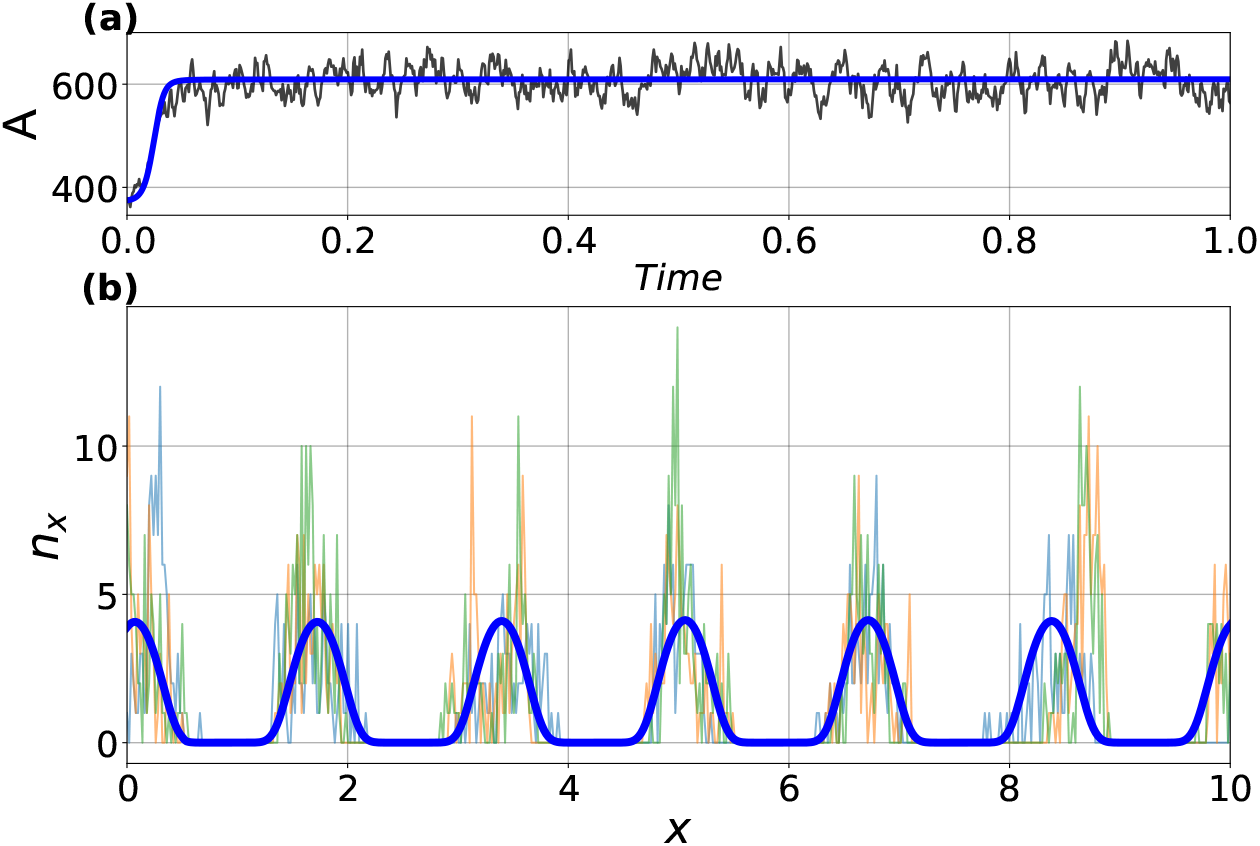
Comparison between the stochastic model and its deterministic limit. A) Population size as a function of time for a single realization of the stochastic process (black line) and the deterministic approximation (blue). B) Spatial distribution of individuals generated by the stochastic dynamics (thin blue, orange, and green lines; each line represents a snapshot of the stationary spatial distribution of individuals) and the deterministic approximation (blue thick curve). For the latter, we used an initial condition *ρ*_+_ + *ϕ*(*x*), where *ϕ*(*x*) is a white noise uncorrelated in space with mean zero and variance *ϵ* ≪ 1, and transformed population density to size by multiplying the value of the density field in each of the PDE integration nodes by the length of the lattice mesh used in the discrete simulations *δx*. Parameters and lattice mesh are the same we used in Figure 2 (e, f). The deterministic simulations run until *t* = 1500, with *dt* = 0.05 and *dx* = 0.008. See Supplementary Material section S6 for details on the algorithm.

To investigate whether organisms aggregate or not, we perform a linear stability analysis of Eq. (2.5). This technique consists in adding a small spatial perturbation to a stable uniform solution of the equation and calculating the perturbation growth rate. If the perturbation growth rate is negative, the uniform solution is stable and patterns do not form. Conversely, the perturbation leads to spatially periodic solutions or patterns if its growth rate is positive (Cross and Hohenberg, 1993). We consider a solution of the form *ρ*(*x, t*) = *ρ*_+_ + *ϵψ*(*x, t*) where *ρ*_+_ is a uniform solution of Eq. (2.5), and *ψ*(*x, t*) an arbitrary perturbation modulated by an amplitude parameter *ϵ* ≪ 1. We insert this solution into Eq. (2.5) and obtain an ordinary differential equation for the dynamics of the perturbation *ψ*(*x, t*). By linearizing and Fourier transforming this differential equation, we obtain the perturbation growth rate as a function of its wavenumber *k* (see Supplementary Material section S2 for details of the calculation). This perturbation growth rate is

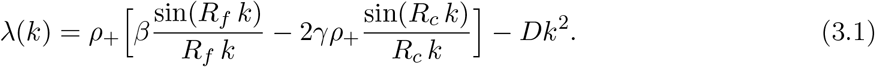

If *λ*(*k*) is positive for a given wavenumber *k*, a perturbation with that wavenumber will grow and create a regular pattern of population density. The wavenumber maximizing *λ*(*k*) in Eq. (3.1), *k*_*max*_, defines the dominant periodicity of the spatial pattern at short times and is related to the periodicity of the long-term spatial pattern of population density. Hence, we can estimate the number of groups *m* that form in a system of size *L* as *m ≈ Lk*_*max*_*/*2*π*. Moreover, we can better understand how the different processes and interactions included in the microscopic model contribute to pattern formation by analyzing term by term all the different contributions to the perturbation growth rate.

First, the linear stability analysis shows that diffusion contributes with a negative term to Eq. (3.1) and hence tends to homogenize population density and eliminate patterns. Second, longrange competition and facilitation enter in the perturbation growth rate via the Fourier transform of their corresponding interaction kernel, which, in the case of the top-hat kernel chosen in our model, are damped oscillatory functions with interaction-specific frequency, magnitude, and sign (Fig. 4). The frequency of each oscillatory function is determined by the interaction range, while the magnitude is determined by the intensity of the intraspecific interaction. The sign preceding the each oscillatory function indicates how competition or facilitation impact population growth, with the negative sign corresponding to competition and the positive one to facilitation.

**Figure 4:**
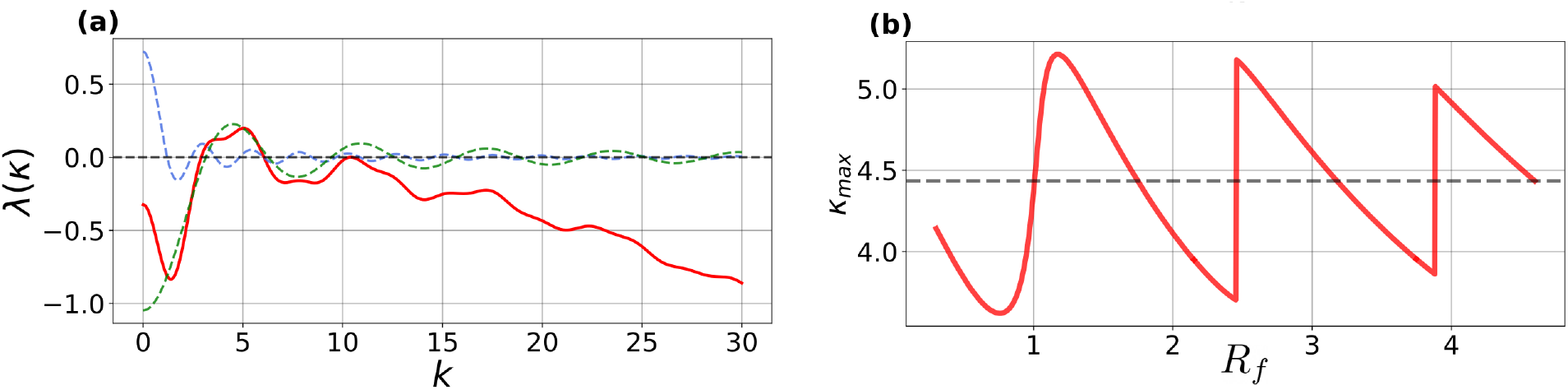
a) Perturbation growth rate as a function of the wavenumber *k* (red). The dashed lines represent the contributions of the facilitation (blue) and competition (green) terms to *λ*(*k*). b) The fastest growing wavenumber, *k*_*max*_, as a function of *R*_*f*_. The grey dashed line is the number of peaks predicted in the absence of facilitation, *β→* 0. We use *r* = *−*2, *D* = 0.001and *β* = 1, *γ* = 1. For panel (a) we choose *R*_*f*_ = 2.6.

To better understand the role of long-range competition and facilitation in the formation of aggregates, we next consider the limit cases in which each of these interactions vanishes or acts on a local scale. In the local competition limit, *R*_*c*_ *→* 0, the perturbation growth rate is always negative because *ρ*_+_ *< β/*(2 *γ*) when populations are uniformly distributed [see Eq. (2.8)]. Therefore, patterns do not form. However, if facilitation is local, *R*_*f*_ *→* 0, or vanishes, *β* = 0, the perturbation growth rate can still be positive for certain wavenumbers, and patterns can potentially form. Varying facilitation makes the fastest-growing wavenumber, and therefore the number of groups, oscillate around the value obtained when long-range interactions are purely competitive. Therefore, longrange competition is a sufficient and necessary condition for pattern formation, and it sets the periodicity of the long-term spatial pattern of population density. Facilitation, on the other hand, plays a secondary role in pattern formation, rearranging the pattern periodicity around the value set by the competition range (Rietkerk and Van de Koppel, 2008). Previous studies have already identified long-range competition as a cause of spatial patterns through the establishment of the so-called exclusion regions, i.e., regions between clusters of organisms in which individuals would compete with individuals from two neighbor groups (Hernández-García and López, 2004; Martínez-García et al., 2013, 2014). In fact, for low diffusion, our simulations show that the distance between aggregates is very close to the competition range *R*_*c*_, as expected when patterns form due to exclusion regions (Hernández-García and López, 2004; Martinez-Garcia et al., 2023; Pigolotti et al., 2007). Moreover, the spatial patterns of population density exhibit aggregates shorter than the range of both non-local interactions, which makes the intensity of competition and facilitation inside an aggregate approximately constant.

### 3.2 The effect of the population spatial distribution on demographic Allee effect

In the previous section, we investigated the conditions in which organisms distribute in non-uniform patterns of population density and quantified the features of the emergent aggregates. Next, we study how aggregation impacts the demographic Allee effect compared to a uniformly distributed population. More specifically, we focus on how group formation affects the main features of a strong demographic Allee effect: the stationary population density, the Allee threshold, and the value of the net growth rate at which extinction is the only stationary state, *r*_*c*_. Because we are interested in the strong Allee effect regime, we limit our analysis to negative density-independent net population growth rates, *r <* 0. In this parameter regime, if the population density is uniform, from Eq. (2.8), we find that *r*_*c*_ = *−β*^2^*/*(4 *γ*) and both *ρ*_+_ and *ρ*_*−*_ exist and are positive for *r* ∈ [*r*_*c*_, 0]. This range of values of *r* defines the region of the parameter space in which the population exhibits a strong demographic Allee effect, with Allee threshold equal to *ρ*_*−*_ and stationary population density *ρ*_+_.

As we already saw from the simulations of the stochastic individual-based dynamics, aggregation increases the stationary population density. The Allee threshold becomes space-dependent, and it is determined by the local density of individuals within the competition and facilitation ranges. This local densities, in turn, depend on the number and spatial arrangement of groups. Finally, *r*_*c*_ decreases due to aggregation (Fig. 5). As a result of these changes in *r*_*c*_ and the Allee threshold, populations exhibiting a self-organized spatial pattern of population density and a component Allee effect can persist in harsher environments and at higher numbers than uniformly distributed populations. Moreover, because spatially structured populations have lower Allee thresholds, they are less susceptible to extinctions caused by environmental perturbations and can recover after extinction following smaller fluctuations than uniformly distributed populations. We obtained these results using the deterministic approximation in Eq. (2.5), which allows us to compute both stable and unstable solutions of our model (see Supplementary Material section S3 for a detailed description of how we obtained the bifurcation diagram in Fig. 5). We further tested these predictions with direct numerical simulations of the individual-level stochastic dynamics and obtained a very good agreement for most values of *r*. The disagreement between the deterministic approximation and stochastic simulations appears for values of *r* close to *r*_*c*_. In this regime, fluctuations in population size can take the population size below the Allee threshold and cause extinctions more easily (Dennis, 2002). Thus, fluctuations become an important driver of population dynamics in this parameter regime, and the mean-field results diverge from the stochastic ones.

**Figure 5:**
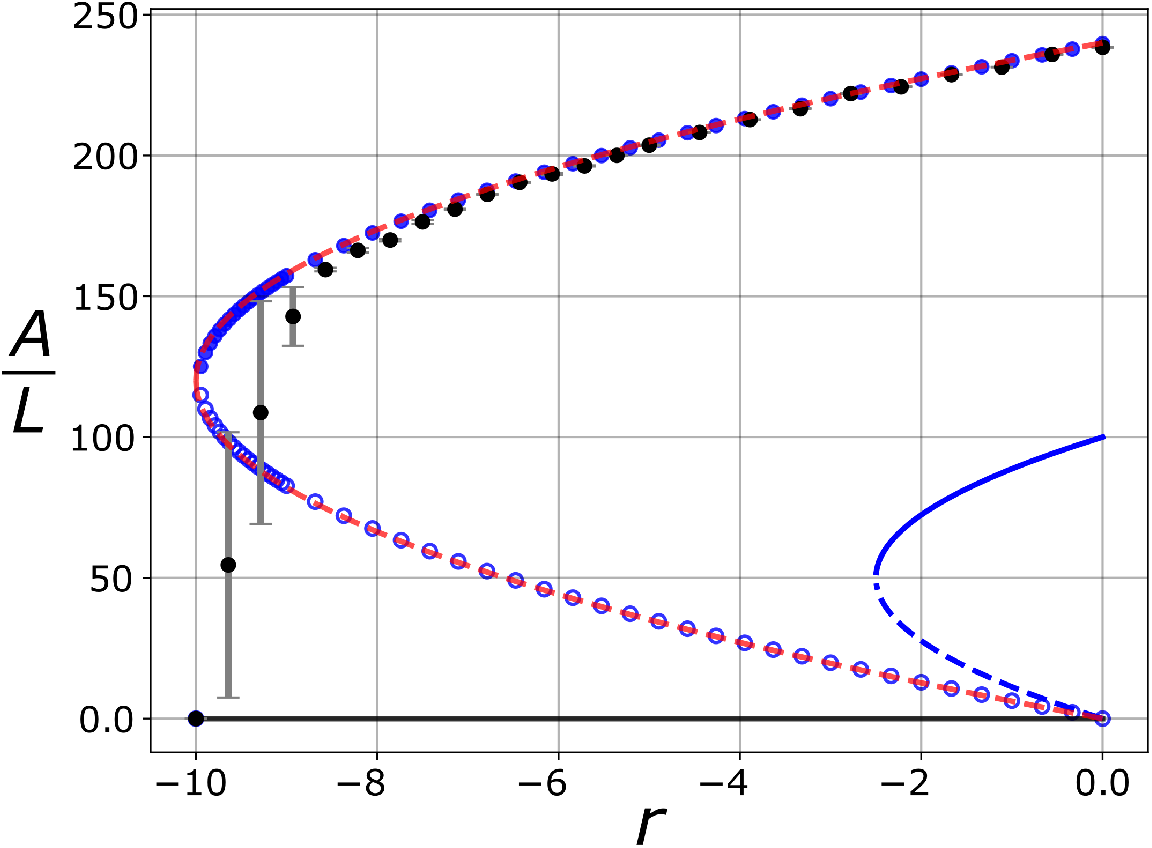
Effect of spatial self-organization on the demographic Allee affect. Population abundance as a function of the net population growth, r, obtained from: the deterministic density equation when patterns develop, Eq. (2.5) (blue points and blue lines); the non-spatial cubic model (blue lines); the meta-population approximation, Eq. (3.3) (dashed red line); and the stochastic dynamics (black circles with error bars indicating the variance of 50 independent realizations). The filled points and the blue solid line represent a stable equilibrium, whereas the empty symbols and blue dashed lines represent unstable equilibrium states. The deterministic simulations run until *t* = 1500, with *dt* = 0.05 and *dx* = 0.008. The stochastic model runs until *t* = 500 with *β* = 10^*−*1^, *γ* = 10^*−*3^, *R*_*f*_ = 0.5, *R*_*c*_ = 1 and *δx* = 0.02. All simulations are done with *L* = 32 and *D* = 10^*−*3^. See Supplementary Material section S6 for details on the numerical methods.

To develop a more mechanistic understanding of how spatial patterns impact the properties of the demographic Allee effect, we further approximate the deterministic equation (2.5) for population density by a network, metapopulation-like description in which each node or population represents a group of individuals and each link represents the existence of inter-group facilitation. We build this approximation based on three features of the spatial patterns of population density. First, all individuals within a group must interact with one another via competition and facilitation. Mathematically, this means that competition and facilitation ranges must be greater than clusters of organisms. Second, individuals of different groups must not compete with each other. In terms of our model, this condition implies that the competition range must be shorter than the distance between pattern aggregates. Finally, if two groups interact with each other via facilitation, this positive interaction must reach all the individuals in both groups. Therefore, the facilitation range must be large enough to encompass all the individuals of a neighbor group. The first two assumptions are only met when diffusion is low, and the spatial structure of the system is determined mainly by the finite-range ecological interactions. The last assumption is correct provided that the first two are met, except for specific values of *R*_*f*_ for which the facilitation range reaches neighboring clusters partially. Considering these three assumptions, the number of individuals in each group changes according to (see Supplementary Material section S4),

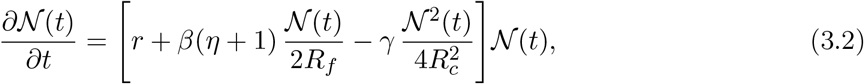

which is a cubic equation for the dynamics of group size, 𝒩. This equation encodes all the information about the underlying network of inter-group interactions in the parameter *η*, which defines the number of groups that interact with a focal group via facilitation, excluding the focal group itself. Solving Eq. (3.2) we can obtain the possible stationary group sizes 𝒩_0_ = 0 (extinction) and:

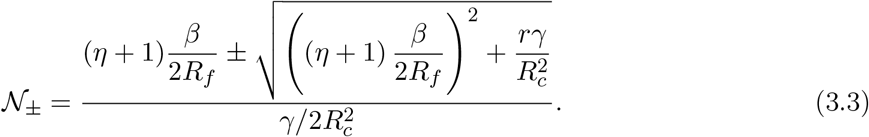

The predictions of this metapopulation-like approximation for *r*_*c*_ and the steady-state population size are in excellent agreement with those of the density equation and the outcome of the stochastic simulations (Fig. 5). In addition, mapping the spatially explicit dynamics to a set of coupled ordinary differential equations allows us to obtain analytical expressions for these two features of the demographic Allee effect in the presence of spatial patterns of population density. The steadystate population size is *A* = *m* 𝒩_+_, where 𝒩_+_ is given by Eq. (3.3) and *m* is the number of groups that we can estimate from the pattern wavelength predicted by the wavenumber that maximizes the perturbation growth rate in Eq. (3.1), *k*_*max*_. Imposing 𝒩_+_ = 𝒩_*−*_ in Eq. (3.3), we can calculate the critical value of the net growth rate that can sustain a non-zero population size,

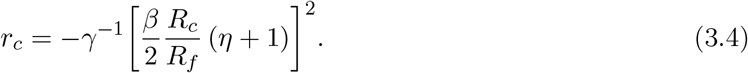

As expected, *r*_*c*_ decreases with increasing facilitation and decreasing competition strength. In addition, *r*_*c*_ decreases when the number of groups that interact with one another increases. More specifically, for certain net growth rates *r*, a population would only be able to survive provided that groups facilitate each other (Fig. 6c), which makes long-range interactions a necessary conditions for population survival. Notice, however, that when the facilitation range increases and groups rely on one another for survival, the whole population becomes less resistant to local perturbations that might cause global extinctions due to the high connectivity between groups.

**Figure 6:**
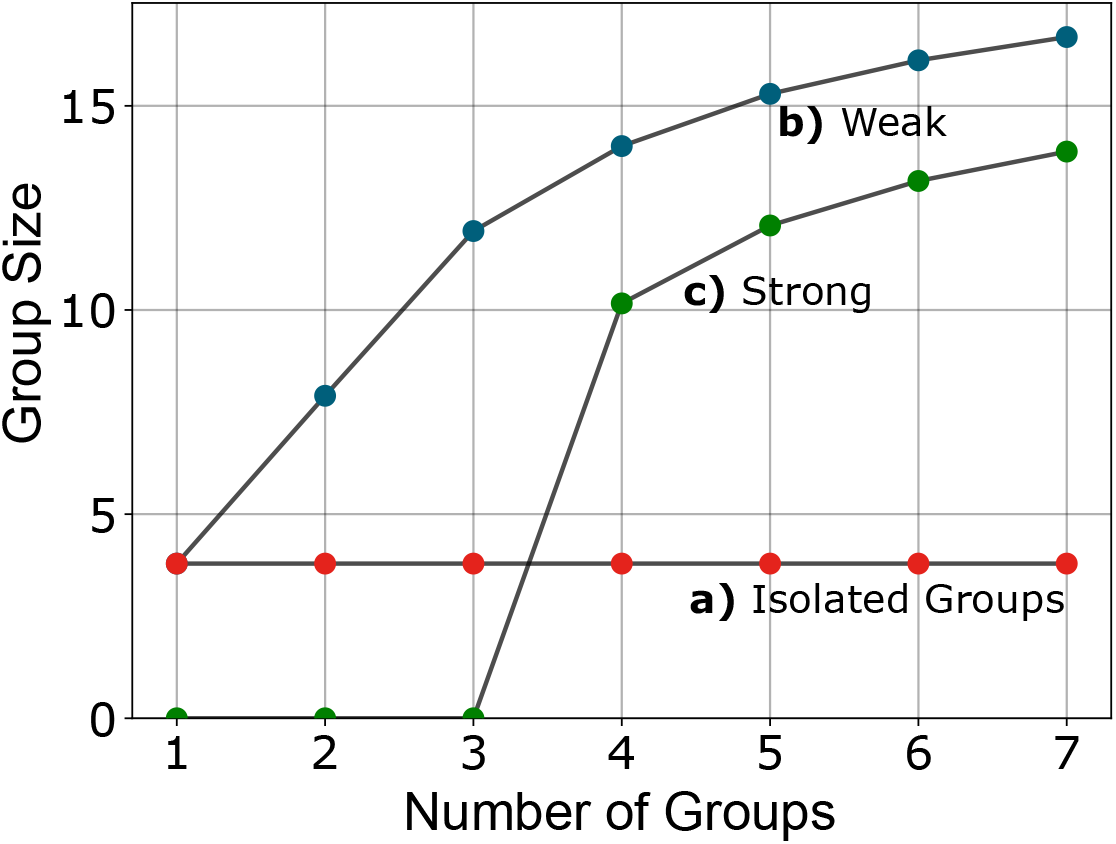
Demographic Allee effect in a population composed of groups. Here, we set the number of groups in the system and compute the size of a single group in the stationary state, 𝒩_+_. The red symbols correspond to a situation in which groups are isolated, 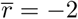 and *Rf* = 0.5; blue and green symbols correspond to cases with inter-group facilitation with *R*_*f*_ = 2 and *r* =*−* 2 (blue) and *r* = *−*100 (green). For all cases, *R*_*c*_ = 1

Organism grouping sets new ways in which the individual-level component Allee effect manifests at the population level and determines the Allee threshold. We analyze these possible outcomes for different numbers of groups and facilitation ranges using the metapopulation-like approximation in Eq. (3.2) that gives the dynamics of each group independently. Mimicking the one-dimensional landscape we used in all previous analyses, we consider that groups are arranged in a line. However, we do not consider periodic boundary conditions to prevent the number of groups from being effectively infinite. If the facilitation range is short so individuals in different groups do not interact with one another, the fitness of the individuals within each aggregate only depends on group size (Fig. 6a), and groups are independent units. In consequence, the formation or extinction of a group does not have any effect on the others, and the minimum population size that ensures population survival is equal to the Allee threshold of one single group, 𝒩_*−*_ from Eq. (3.3) with *η* = 0. If the facilitation range is such that groups interact with one another, the fitness of the individuals can increase significantly due to the presence of neighbor groups. As a consequence, group size increases in the presence of more groups (Fig. 6b and 6c), and the Allee threshold is 𝒩_*−*_ from Eq. (3.3) with *η >* 0. For very harsh environmental conditions (low *r*) the population only survives if groups facilitate one another (Fig. 6c).

Finally, we computed the stationary-state population size as a function of the diffusion coefficient to evaluate the range of diffusion intensity at which the first two assumptions underlying the group-level approximation in Eq. (3.2) remain valid (Fig. 7a). Consistently with the simulations of the stochastic individual-based dynamics (Fig. 2), we observe that the total population abundance decreases as diffusion increases. In the low-diffusion regime, the population abundance agrees with the predictions of the meta-population approximation. However, as diffusion increases, diffusion takes control of the spatial dynamics, and the assumptions underlying the metapopulation approximation stop being valid. As a result, the population density decreases until diffusion reaches a critical value (black dashed line in Figure 7a), at which patterns do not form and the population abundance is equal to that predicted by models assuming uniformly distributed individuals. We also observe this decrease in population density in the spatial patterns of population density, which tend to become uniform as diffusion increases (Fig. 7b).

**Figure 7:**
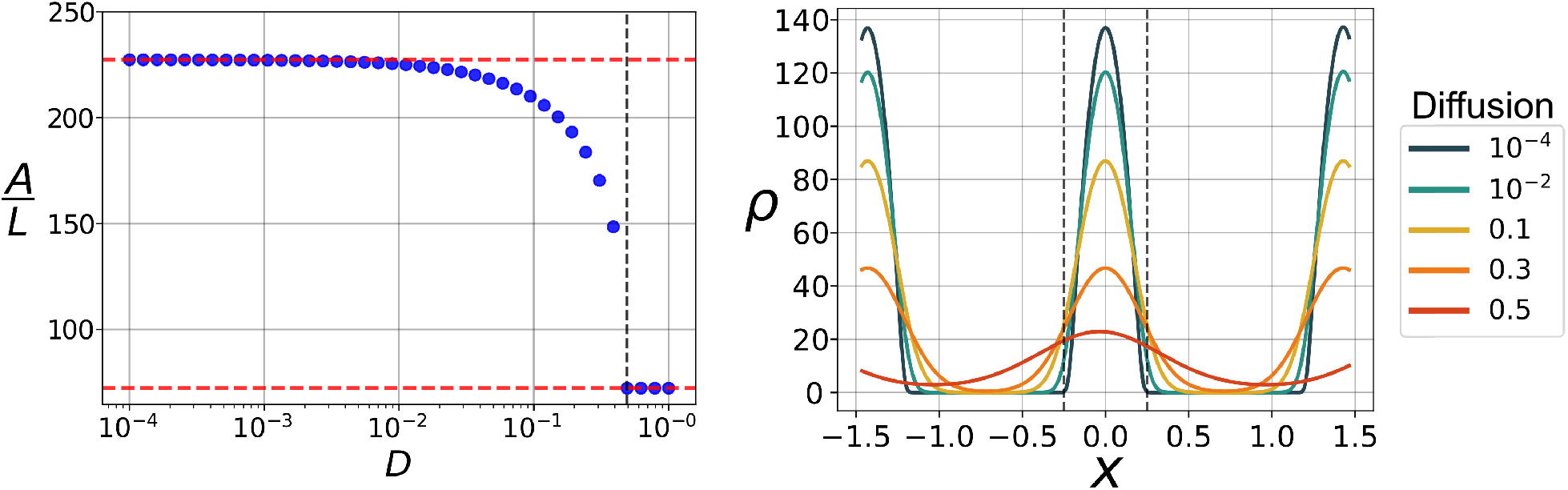
(a) Effect of increasing diffusion on the population abundance. Self-organized spatial patterns disappear when diffusion increases and the population abundance decreases from the metapopulation prediction 𝒩_+_ to the uniform solution *ρ*_+_. (b) Effect of diffusion on spatial patterns, stationary patterns of population density for different diffusion intensities (color code indicated in the legend). The black dashed lines limit the extent of the facilitation range, *R*_*f*_. Parameter values (for both panels): *t* = 2 *×* 10^4^, with *dt* = 0.05, *dx* = 0.008, *L* = 32 and parameters: *r* = *−*2, *R*_*f*_ = 0.5 and *R*_*c*_ = 1.

## 4 Discussion

We theoretically investigated the demographic consequences of a component Allee effect across various levels of spatial organization (Fig. 1). Our framework incorporates a component Allee effect arising from reproductive facilitation, which makes the reproduction rates of a focal individual increase with the population density within its neighborhood, and growth limitation caused by intraspecific competition. Extending our analysis to other types of individual-level interactions leading to component Allee effects, such as social behaviors, mate limitation, or environmental conditioning (Courchamp et al., 2008; Oro, 2020b) is straightforward. We focused on quantifying the impact of the spatial distribution of organisms on specific features of the demographic Allee effect, such as the Allee threshold, the long-term total population size, and the lowest value of the density-independent growth rate for which the population survives. We measured these quantities in both uniformly and non-uniformly distributed populations.

Our approach, similarly to Surendran et al. (2020), differs from non-spatial and spatially implicit metapopulation models by explicitly considering the range of interaction for both reproductive facilitation and crowding effects. This level of detail is partially captured by metapopulation models, which assume that individuals only interact with others within the same population. Metapopulation frameworks, however, assume a fixed population structure in groups, whereas groups emerges naturally from individual-level processes in our model. This explicit description of the processes the lead to grouping allows us to identify the individual-level processes that control for each of the population-level features of the demographic Allee effect and subsequently manipulate them to understand how different spatial structures impact the demographic Allee effect.

In addition, limiting the mechanisms responsible for the component Allee effect to a finite neighborhood around each focal individual makes the population dynamics and the features of the emergent demographic Allee effect depend on local, instead of total, population densities. For example, the Allee threshold becomes a local feature of the population that depends on the density of individuals within a given region of the landscape. This locality of the Allee threshold might enable the survival of local populations in situations where the global density is very low, which is especially relevant when spatial fluctuations in population density are high, such as in the presence of clumps of organisms or groups. This strong dependence of the Allee threshold on the spatial population structure might help to explain field studies reporting population survival at low global population densities (Lundquist and Botsford, 2011; Rijnsdorp and Vingerhoed, 2001; Woodroffe, 2011).

Our model also provides the appropriate theoretical framework to formalize the group Allee effect and integrate it within a unifying modeling approach (Angulo et al., 2018, 2013). When organisms aggregate, one can consider the groups as the fundamental units of the population. If competition acts on a longer range than facilitation, these groups are independent units that do not interact with one another. In consequence, the component Allee effect impacts the demographics of a single group, resulting in a demographic group Allee effect that only determines the population dynamics within that group. This same argument can be extended to cases in which facilitation acts on a longer range than competition. In this limit, groups interact with one another, which can result in a group-level Allee effect when the fitness of a group increases in the presence of neighbors. This group-level component Allee effect scales to the population level by creating an emergent demographic Allee effect acting on groups that can even result in the existence of a minimum number of groups to ensure population survival.

Beyond group-level processes, spatial heterogeneities in population density favor population survival as long as the density within a region of the landscape is locally above the Allee threshold. Moreover, because groups in our model form in response to long-range competition, aggregation minimizes competition and results in larger global population sizes that are less prone to extinction due to demographic fluctuations (Dennis, 2002). Aggregation also lowers the Allee threshold significantly, which favors the persistence of local populations at lower population densities. This local decrease in the Allee threshold is different from the effective decrease in the global Allee threshold discussed above, which is related to the locality of the Allee threshold rather than to its value. Finally, as found in previous studies, our model predicts that aggregated populations can survive in harsher environments than uniformly distributed populations. That is, uniformly distributed populations exhibit a higher value of *r*_*c*_ than populations that develop self-organized spatial patterns (Surendran et al., 2020).

Our model provides the simplest framework to study Allee effects across levels of spatial organization and a unifying theoretical approach to understand how Allee effects operate for different population structures. To keep it as simple as possible, we made some simplifying model assumptions. The choice of the component Allee effect, as we discussed before, can be easily changed by modifying the set of individual-level demographic reactions. Other assumptions, such as the choice of the interaction kernels, would not change our results provided that they lead to spatial pattern formation (Colombo et al., 2023; Martínez-García et al., 2013; Pigolotti et al., 2007). One could also consider a different mechanism responsible for spatial pattern formation, and our results would hold provided that spatial patterns emerge in the form of clumps of population density. We considered non-local interactions as the pattern-forming mechanism because it is the most straight-forward way to create aggregation patterns (Martínez-García et al., 2014). An interesting direction for future research, however, would be to consider alternative pattern-forming interactions, such as density-dependent movement or resource-consumer interactions, leading to a larger variety of spatial patterns in population density, such as labyrinths and gaps (Liu et al., 2013; Martinez-Garcia et al., 2015, 2022; Rao and Kang, 2016; Rietkerk and Van de Koppel, 2008). Finally, our modeling framework is also easily extendable to include interactions between several species (Maciel and Martinez-Garcia, 2021; Simoy and Kuperman, 2023), thus providing a theoretical tool to investigate community-level consequences of different component Allee effects.

## 5 Conclusions

We investigated the demographic consequences of an individual-level component Allee effect in a spatially extended population (Fig. 1). We departed from a mechanistic description of how the vital rates of a focal individual depend on the density of conspecifics around it. Our model, therefore, accounts for spatial processes both through the spatial population structure and the range of the different interactions among them. We considered the most straightforward set of processes leading to a demographic Allee effect, which in the non-spatial limit collapses to a cubic model (Kot, 2001; Méndez et al., 2019). Starting from this description of the individual vital rates, we present a series of mathematical techniques to investigate the population-level how a component Allee effect manifests across various characteristic spatial scales of the population.

For the specific component Allee effect we studied here, we show that aggregation changes three main population-level features characteristics of Allee effects. First, aggregation enhances population density locally and thus allows the population to persist in harsh environments where uniformly distributed individuals would go extinct. Second, aggregation results in localized subpopulations that follow independent dynamics from one another and might eliminate the population-level Allee effect. Finally, aggregation decreases competition by limiting its effect to individuals within the same group. Consequently, aggregation reduces the Allee threshold and increases the total population size. More generally, our work emphasizes the potential that models developed from a rigorous description of the individual-level interactions and processes have to improve our understanding of observed patterns and trends in population dynamics.

## Acknowledgments

This work was partially funded by the Center of Advanced Systems Understanding (CASUS), which is financed by Germany’s Federal Ministry of Education and Research (BMBF) and by the Saxon Ministry for Science, Culture and Tourism (SMWK) with tax funds on the basis of the budget approved by the Saxon State Parliament. This work was also funded by FAPESP through a Master Fellowship no. 2020/15643-8 (D.C.P.J), a BIOTA Young Investigator Research Grant no. 2019/05523-8 (D.C.P.J. and R.M.-G.), and ICTP-SAIFR grant no. 2016/01343-7; the Abdus Salam ICTP through the Associate’s Programme, and the Simons Foundation through grant no. 284558FY19.

## References

Allee, W. and Bowen, E. S. (1932). Studies in animal aggregations: mass protection against colloidal silver among goldfishes. Journal of Experimental Zoology, 61(2):185–207.

Allee, W. and Rosenthal, G. (1949). Group survival value for philodina rosola, a rotifer. Ecology, 30(3):395–397.

Allee, W. C. (1938). Social life of animals. Number Edn 1. William Heineman Ltd, London and Toronto.

Allee, W. C., Park, O., Emerson, A. E., Park, T., Schmidt, K. P., et al. (1949). Principles of Animal Ecology. Number Edn 1. WB Saundere Co. Ltd.

Angulo, E., Luque, G. M., Gregory, S. D., Wenzel, J. W., Bessa-Gomes, C., Berec, L., and Courchamp, F. (2018). Allee effects in social species. Journal of Animal Ecology, 87(1):47–58.

Angulo, E., Rasmussen, G. S., Macdonald, D. W., and Courchamp, F. (2013). Do social groups prevent Allee effect related extinctions?: The case of wild dogs. Frontiers in zoology, 10(1):1–14.

Ashman, T.-L., Knight, T. M., Steets, J. A., Amarasekare, P., Burd, M., Campbell, D. R., Dudash, M. R., Johnston, M. O., Mazer, S. J., Mitchell, R. J., et al. (2004). Pollen limitation of plant reproduction: ecological and evolutionary causes and consequences. Ecology, 85(9):2408–2421.

Asmussen, M. A. (1979). Density-dependent selection ii. the Allee effect. The American Naturalist, 114(6):796–809.

Colombo, E. H., López, C., and Hernández-García, E. (2023). Pulsed interaction signals as a route to biological pattern formation. Physical Review Letters, 130(5):058401.

Courchamp, F., Berec, L., and Gascoigne, J. (2008). Allee effects in ecology and conservation. OUP Oxford.

Crews, D., Grassman, M., and Lindzey, J. (1986). Behavioral facilitation of reproduction in sexual and unisexual whiptail lizards. Proceedings of the National Academy of Sciences of the United States of America, 83(24):9547–9550.

Cross, M. C. and Hohenberg, P. (1993). Pattern formation outside of equilibrium. Reviews of Modern Physics, 65(3).

Cushing, J. (1988). The Allee effect in age-structured population dynamics. In Hallam, T. G., Gross, L., and Levin, S., editors, Mathematical Ecology-Proceedings Of The Autumn Course Research Seminars International Ctr For Theoretical Physics, pages 479–505. World Scientific Publ.

Dechmann, D. K., Kranstauber, B., Gibbs, D., and Wikelski, M. (2010). Group hunting—a reason for sociality in molossid bats? PLoS one, 5(2):e9012.

Dennis, B. (1981). Extinction and waiting times in birth-death processes: applications to endangered species and insect pest control. Statistical distributions in scientific work, 6:289–301.

Dennis, B. (1989). Allee effects: population growth, critical density, and the chance of extinction. Natural Resource Modeling, 3(4):481–538.

Dennis, B. (2002). Allee effects in stochastic populations. Oikos, 96(3):389–401.

Doi, M. (1976). Stochastic theory of diffusion-controlled reaction. Journal of Physics A: Mathematical and General, 9(9):1479.

Drake, J. and Kramer, A. (2011). Allee effects. Nature Education Knowledge, 3(10):2.

Garrett, K. and Bowden, R. (2002). An Allee effect reduces the invasive potential of tilletia indica. Phytopathology, 92(11):1152–1159.

Ghazoul, J. (2005). Buzziness as usual? Questioning the global pollination crisis. Trends in ecology & evolution, 20(7):367–373.

Gillespie, D. T. (1977). Exact stochastic simulation of coupled chemical reactions. The journal of physical chemistry, 93555(1):2340–2361.

Guy, C., Smyth, D., and Roberts, D. (2019). The importance of population density and interindividual distance in conserving the european oyster ostrea edulis. Journal of the Marine Biological Association of the United Kingdom, 99(3):587–593.

Hernández-García, E. and López, C. (2004). Clustering, advection, and patterns in a model of population dynamics with neighborhood-dependent rates. Physical Review E, 70(1):016216.

Hsu, P.-H. and Fredrickson, A. (1975). Population-changing processes and the dynamics of sexual populations. Mathematical Biosciences, 26(1-2):55–78.

Kanarek, A. R., Webb, C. T., Barfield, M., and Holt, R. D. (2013). Allee effects, aggregation, and invasion success. Theoretical ecology, 6(2):153–164.

Keitt, T. H., Lewis, M. A., and Holt, R. D. (2001). Allee effects, invasion pinning, and species’ borders. The American Naturalist, 157(2):203–216.

Kostitzin, V. (1940). Sur la loi logistique et ses généralisations. Acta Biotheoretica, 5(3):155–159.

Kot, M. (2001). Harvest models: bifurcations and breakpoints. In Elements of Mathematical Ecology, pages 13–25. Cambridge University Press.

Kramer, A. M., Dennis, B., Liebhold, A. M., and Drake, J. M. (2009). The evidence for Allee effects. Population Ecology, 51(3):341–354.

Lande, R. (1987). Extinction thresholds in demographic models of territorial populations. The American Naturalist, 130(4):624–635.

Le Cadre, S., Tully, T., Mazer, S. J., Ferdy, J.-B., Moret, J., and Machon, N. (2008). Allee effects within small populations of Aconitum napellus ssp. lusitanicum, a protected subspecies in Northern France. New Phytologist, 179(4):1171–1182.

Lerch, B. A., Nolting, B. C., and Abbott, K. C. (2018). Why are demographic Allee effects so rarely seen in social animals? Journal of Animal Ecology, 87(6):1547–1559.

Levitan, D. R. (2005). The Allee effect in the sea. Marine Conservation Biology: the science of maintaining the sea’s biodiversity, pages 47–57.

Liermann, M. and Hilborn, R. (2001). Depensation: evidence, models and implications. Fish and Fisheries, 2(1):33–58.

Liu, Q.-X., Doelman, A., Rottschäfer, V., de Jager, M., Herman, P. M., Rietkerk, M., and van de Koppel, J. (2013). Phase separation explains a new class of self-organized spatial patterns in ecological systems. Proceedings of the National Academy of Sciences, 110(29):11905–11910.

Lundquist, C. J. and Botsford, L. W. (2011). Estimating larval production of a broadcast spawner: the influence of density, aggregation, and the fertilization Allee effect. Canadian Journal of Fisheries and Aquatic Sciences, 68(1):30–42.

Luque, G. M., Giraud, T., and Courchamp, F. (2013). Allee effects in ants. Journal of Animal Ecology, 82(5):956–965.

Luzuriaga, A. L., Escudero, A., Albert, M. J., and Giménez-Benavides, L. (2006). Population structure effect on reproduction of a rare plant: beyond population size effect. Botany, 84(9):1371–1379.

Maciel, G. A. and Lutscher, F. (2015). Allee effects and population spread in patchy landscapes. Journal of Biological Dynamics, 9(1):109–123.

Maciel, G. A. and Martinez-Garcia, R. (2021). Enhanced species coexistence in Lotka-Volterra competition models due to nonlocal interactions. Journal of Theoretical Biology, 530:110872.

Martinez-Garcia, R., Cabal, C., Calabrese, J. M., Hernández-García, E., Tarnita, C. E., López, C., and Bonachela, J. A. (2023). Integrating theory and experiments to link local mechanisms and ecosystem-level consequences of vegetation patterns in drylands. Chaos, Solitons & Fractals, 166:112881.

Martínez-García, R., Calabrese, J. M., Hernández-García, E., and López, C. (2013). Vegetation pattern formation in semiarid systems without facilitative mechanisms. Geophysical Research Letters, 40(23):6143–6147.

Martínez-García, R., Calabrese, J. M., Hernández-García, E., and López, C. (2014). Minimal mechanisms for vegetation patterns in semiarid regions. Philosophical Transactions of the Royal Society A: Mathematical, Physical and Engineering Sciences, 372(2027):20140068.

Martinez-Garcia, R., Murgui, C., Hernández-García, E., and López, C. (2015). Pattern Formation in Populations with Density-Dependent Movement and Two Interaction Scales. PLoS ONE, 10:e0132261.

Martinez-Garcia, R., Tarnita, C. E., and Bonachela, J. A. (2022). Self-organized patterns in ecological systems: from microbial colonies to landscapes. Emerging Topics in Life Sciences, 6(3):245–258.

Méndez, V., Assaf, M., Masó-Puigdellosas, A., Campos, D., and Horsthemke, W. (2019). Demographic stochasticity and extinction in populations with Allee effect. Physical Review E, 99(2):022101.

Nowak, K. and Lee, P. C. (2011). Demographic structure of zanzibar red colobus populations in unprotected coral rag and mangrove forests. International Journal of Primatology, 32(1):24–45.

Oro, D. (2020a). Extinction, nonlinear dynamics, and sociality. In Oro (2020b), pages 114–127.

Oro, D. (2020b). Perturbation, behavioural feedbacks, and population dynamics in social animals: when to leave and where to go. Oxford University Press, USA.

Orr, H. A. (2009). Fitness and its role in evolutionary genetics. Nature Reviews Genetics, 10(8):531–539.

Padrón, V. and Trevisan, M. C. (2000). Effect of aggregating behavior on population recovery on a set of habitat islands. Mathematical biosciences, 165(1):63–78.

Peliti, L. (1985). Path integral approach to birth-death processes on a lattice. Journal de Physique, 46(9):1469–1483.

Pigolotti, S., López, C., and Hernández-García, E. (2007). Species clustering in competitive lotka-volterra models. Physical review letters, 98(25):258101.

Rao, F. and Kang, Y. (2016). The complex dynamics of a diffusive prey–predator model with an Allee effect in prey. Ecological complexity, 28:123–144.

Rietkerk, M. and Van de Koppel, J. (2008). Regular pattern formation in real ecosystems. Trends in ecology & evolution, 23(3):169–175.

Rijnsdorp, A. and Vingerhoed, B. v. (2001). Feeding of plaice pleuronectes platessa l. and sole solea solea (l.) in relation to the effects of bottom trawling. Journal of Sea Research, 45(3-4):219–229.

Silliman, B. R., Schrack, E., He, Q., Cope, R., Santoni, A., van der Heide, T., Jacobi, R., Jacobi, M., and van de Koppel, J. (2015). Facilitation shifts paradigms and can amplify coastal restoration efforts. Proceedings of the National Academy of Sciences, 112(46):14295–14300.

Simoy, M. I. and Kuperman, M. N. (2023). Non-local interaction effects in models of interacting populations. Chaos, Solitons & Fractals, 167:112993.

Snaith, T. V. and Chapman, C. A. (2008). Red colobus monkeys display alternative behavioral responses to the costs of scramble competition. Behavioral Ecology, 19(6):1289–1296.

Stephens, P. A., Frey-roos, F., Arnold, W., and Sutherland, W. J. (2002). Model complexity and population predictions. the alpine marmot as a case study. Journal of Animal Ecology, 71(2):343–361.

Stephens, P. A., Sutherland, W. J., and Freckleton, R. P. (1999). What is the Allee effect? Oikos, xpages 185–190.

Sun, G.-Q. (2016). Mathematical modeling of population dynamics with Allee effect. Nonlinear Dynamics, 85(1):1–12.

Surendran, A., Plank, M. J., and Simpson, M. J. (2020). Population dynamics with spatial structure and an Allee effect. Proceedings of the Royal Society A: Mathematical, Physical and Engineering Sciences, 476:1–19.

Tammes, P., Klomp, H., and Van Montfort, M. (1964). Sexual reproduction and underpopulation. Archives Néerlandaises de Zoologie, 16(1):105–110.

Täuber, U. C. (2007). Field-Theory Approaches to Nonequilibrium Dynamics, pages 295–348. Springer Berlin Heidelberg, Berlin, Heidelberg.

Tcheslavskaia, K., Brewster, C. C., and Sharov, A. A. (2002). Mating success of gypsy moth (lepidoptera: Lymantriidae) females in southern wisconsin. The Great Lakes Entomologist, 35(1):1.

Thomas, J. D. and Benjamin, M. (1974). The Effects of Population Density on Growth and Reproduction of Biomphalaria glabrata (Say) (Gasteropoda: Pulmonata). The Journal of Animal Ecology, 43(1):31.

Volterra, V. (1938). Population growth, equilibria, and extinction under specified breeding conditions: a development and extension of the theory of the logistic curve. Human Biology, 10(1):1–11.

Wagenius, S. (2006). Scale dependence of reproductive failure in fragmented echinacea populations. Ecology, 87(4):931–941.

Wang, Y., Shi, J., and Wang, J. (2019). Persistence and extinction of population in reaction– diffusion–advection model with strong Allee effect growth. Journal of mathematical biology, 78(7):2093–2140.

Woodroffe, R. (2011). Demography of a recovering african wild dog (lycaon pictus) population. Journal of Mammalogy, 92(2):305–315.

Woodroffe, R., O’Neill, H. M., and Rabaiotti, D. (2020). Within-and between-group dynamics in an obligate cooperative breeder. Journal of Animal Ecology, 89(2):530–540.

