## Supplementary Information for "Demographic effects of aggregation in the presence of a component Allee effect"

### Supplementary Material for: 'Demographic effects of aggregation in the presence of a component Allee effect'

**S2 Linear stability analysis of Eq. 2.5** **7**

**S3 Numerical computation of the stationary states of Eq. 2.5** **8**

**S4 Derivation of the group-level approximation in Eq. 3.2** **8**

**S5 Model nondimensionalization** **9**

**S6 Numerical methods** **10**

S6.1 Stochastic individual-based simulations . . . . . . . . . . . . . . . . . . . . . . . . . . . 10

S6.2 Integration of the population-level approximation, Eq. 2.5 . . . . . . . . . . . . . . . . 11

**S7 Supplementary Figure for the two-dimensional case** **12**

#### S1 Derivation of the population-level approximation Eq. 2.5

In this section, we detail the steps to obtain a deterministic equation for the dynamics of the density of individuals starting from the stochastic individual-level reactions. To this end, we apply the Doi-Peliti formalism, which is a field-theoretical approach, developed in the context of statistical field theory, that uses path integrals to map an individual-level stochastic dynamics to a continuum description in terms of a density field (Doi, 1976; Hernández-García and López, 2004; Peliti, 1985; Täuber, 2007).

##### S1.1 Derivation of the Master equation

The Master equation characterizes the evolution of the probability that a system following a stochastic dynamics is in a specific state at a given time $t$. In our case, the Master equation describes how the probability of finding a certain population size and spatial distribution of individuals across the lattice nodes changes with time. We denote this lattice configuration by a vector $\eta$ that specifies the number of individuals in each lattice node: $\eta = \{...n_{i-1},\, n_i,\, n_{i+1}...\}$. Each lattice node coordinate, labeled by the index $i$, can be mapped to a spatial coordinate $x_i$ using the transformation $x_i = i\,\delta x$, where $\delta x$ is the distance between two adjacent lattice nodes.

To construct the Master equation, we need to obtain the global transition rates, $\Omega(\eta \to \eta')$, that define the probabilistic transitions between two lattice configurations $\eta$ and $\eta'$ and will depend on the birth, death, and movement stochastic events. Using these global transition rates, we can write the Master equation as

$$\frac{\partial P(\eta,t)}{\partial t} = \sum_{\eta'} \Omega(\eta' \to \eta) P(\eta',t) - \Omega(\eta \to \eta') P(\eta,t) \tag{S1.1}$$

###### Contribution of birth processes to the global transition rates

Birth processes contribute to the appearance of a new individual in a focal lattice position $x$ via density-independent reproduction and facilitation. These processes are represented by the following biological reactions

$$\bigcirc_x \xrightarrow{b} \bigcirc_x + \bigcirc_x, \tag{S1.2}$$

$$\bigcirc_x + \bigcirc_{x'} \xrightarrow{\frac{\beta}{2R_f}} \bigcirc_x + \bigcirc_{x'} + \bigcirc_x, \tag{S1.3}$$

where the reaction (S1.3) only takes place if $|x - x'| \leq R_f$. We can decompose the global transition rate resulting from these birth processes in two demographic rates $W$, one corresponding to each of the reactions that can potentially contribute to the transition from a configuration $\eta$ to $\eta' = \{n_x + 1\}_\eta$:

$$\Omega\big(\eta \to \{n_x + 1\}_\eta\big) = W_b(n_x) + W_\beta(\eta, x), \tag{S1.4}$$

where $\{n_x + 1\}_\eta$ denotes a lattice configuration in which all nodes have the same number of individuals as in the configuration $\eta$ except the node with spatial coordinate $x$, where the occupancy has increased in one unity. $W_b(n_x)$ and $W_\beta(\eta, x)$ are the demographic rates in which the reactions for density-independent birth and facilitation, (S1.2) and (S1.3) respectively, generate individuals at the position $x$.

The demographic rate corresponding to density-independent birth increases linearly with the number of individuals in the position $x$

$$W_b(n_x) = b\, n_x. \tag{S1.5}$$

For the facilitation demographic rate, however, we need to take into account the long-range of the interaction and thus the fact that reproduction at the lattice coordinate $x$ depends on the number of individuals within the lattice nodes such that $|x - x'| \leq R_f$. For example, the demographic rate in which the facilitation reaction (S1.3) generates an individual in $x$ changes depending on whether $x'$ is equal or different from $x$. This difference exists because pairwise facilitation only increases the number of individuals at $x$ half of the times if $x' \neq x$, leading to a new individual at $x'$ the other half. For $x = x'$, however, the new individual is always located at $x$. Thus, considering both the number of pairs we can form for $x = x'$ and $x \neq x'$, the facilitation demographic rate is

$$W_\beta(\eta, x) = \frac{\beta}{2R_f}\left[\overbrace{\binom{n_x}{2}}^{x'=x} + \underbrace{\frac{1}{2}(N_x^b - n_x)n_x}_{x'\neq x}\right] \tag{S1.6}$$

which simplifies to

$$W_\beta(\eta, x) = \frac{\beta\, n_x}{4\, R_f}\left(N_x^b - 1\right). \tag{S1.7}$$

##### Contribution of death processes to the demographic transition rates

Individuals die in each lattice coordinate $x$ due to density-independent death and competition, which we can write in terms of biological reactions as

$$\bigcirc_x \xrightarrow{d} \emptyset, \tag{S1.8}$$

$$\bigcirc_x + \bigcirc_{x'} + \bigcirc_{x''} \xrightarrow{\frac{\gamma}{4R_c^2}} \bigcirc_{x'} + \bigcirc_{x''}, \tag{S1.9}$$

(S1.9) only takes place if both pairwise distances $|x - x'| \leq R_c$ and $|x - x''| \leq R_c$. The global transition rate due to death processes is

$$\Omega(\eta \to \{n_x - 1\}_\eta) = W_d(n_x) + W_\gamma(\eta, x) \tag{S1.10}$$

where $W_d(n_x)$ and $W_\gamma(\eta, x)$ are the demographic rates in which the density-independent death and competition reactions, (S1.8) and (S1.9) respectively, eliminate an individual at lattice coordinate $x$.

Similarly to the density-independent birth process, the density-independent death demographic rate increases linearly with $n_x$

$$W_d(n_x) = d\, n_x, \tag{S1.11}$$

and extending the same combinatorial arguments used to derive Eq. (S1.7) we obtain the ternary competition demographic rate that results in the death of an individual at lattice coordinate $x$,

$$W_\gamma(\eta, x) = \frac{\gamma}{6} \frac{n_x}{4R_c^2} \left(N_x^d - 1\right) \left(N_x^d - 2\right) \tag{S1.12}$$

**Contribution of movement to the demographic transition rates**

The stochastic event that leads to individual movement is

$$\bigcirc_x \xrightarrow{h} \bigcirc_{x \pm \delta x}. \tag{S1.13}$$

where the subscript indicates the location of the individual in spatial coordinates. Because this is a density-independent process, the global rate is linear in $n_x$ and is given by

$$\Omega(\eta \to \{n_x - 1, n_{x'} + 1\}_\eta) = h n_x. \tag{S1.14}$$

where $x' = x + \delta x$ is a nearest neighbor of $x$.

Finally, combining Eqs. (S1.4), (S1.10), and (S1.14) for the global transition rates and scaling $\beta \to 2\beta$ and $\gamma \to 2\gamma$, we can obtain the explicit form of the Master equation. Here we map the positions to their lattice site counterparts $x \mapsto i$.

$$\begin{aligned}
\frac{\partial P(\eta, t)}{\partial t} \;=\;& b \sum_i (n_i - 1) P\left(\{n_i - 1\}_\eta, t\right) - n_i P(\eta, t) + d \sum_i (n_i + 1) P\left(\{n_i + 1\}_\eta, t\right) - n_i P(\eta, t) \\
&+ \frac{\beta}{2R_f} \sum_i (n_i - 1)(N_i^f - 2) P\left(\{n_i - 1\}_\eta, t\right) - n_i (N_i^f - 1) P(\eta, t) \\
&+ \frac{\gamma}{4R_c^2} \sum_i (n_i + 1) N_i^c (N_i^c - 1) P\left(\{n_i + 1\}_\eta, t\right) - n_i (N_i^c - 1)(N_i^c - 2) P\left(\eta, t\right) \\
&+ h \sum_{\langle ij \rangle} (n_i + 1) P\left(\{n_i + 1,\, n_j - 1\}_\eta, t\right) - n_i P(\eta, t).
\end{aligned} \tag{S1.15}$$

where $n_i$ is the number of individuals at the lattice site $i$ and $N_i^\alpha$ denotes the number of individuals within the range $R_\alpha$ centered at $i$. The notation $\langle ij \rangle$ specifies that the sum is performed over the nearest neighbors of $i$.

#### S1.2 Description of the population configuration in terms of a Fock space

Individuals in our discrete model are represented by indistinguishable particles distributed within the cells of a one-dimensional lattice. Therefore, as introduced in Section S1.1, we can describe the state of the system at a given time if we know the number of individuals (particles) in each lattice cell (lattice configuration), $\eta = \{n_0...n_i,\ n_{i+1},\ n_{i+2}...n_N\}$. To apply the Doi-Peliti formalism, we first need to define a Fock space for each lattice node with a basis given by the occupancy number basis and their corresponding creation and annihilation operators. These operators will be responsible for updating the state of each lattice node by changing the number of particles they contain. Therefore, these creation and annihilation operators are related to the demographic rates in our model. Using Dirac notation, the occupancy number basis is a set of vectors $|n_i\rangle$, giving the number of individuals, $n$, in each lattice cell $i$. The creation and annihilation operators, $a_i^\dagger$ and $a_i$, respectively, act on each vector of this basis following

$$a_i^\dagger |n_i\rangle = |n_i + 1\rangle \tag{S1.16}$$

$$a_i |n_i\rangle = n_i |n_i - 1\rangle \tag{S1.17}$$

and have bosonic commutation relations given by

$$\left[a_i, a_j^\dagger\right] = a_i a_j^\dagger - a_j^\dagger a_i = \delta_{ij} \tag{S1.18}$$

$$\left[a_i^\dagger, a_j^\dagger\right] = [a_i, a_j] = 0. \tag{S1.19}$$

Finally, using the occupancy number basis for each lattice node and Dirac's notation, we can express the basis associated with the lattice configuration $\eta$ as the tensor product of the basis of each site

$$|\eta\rangle = |n_0\rangle \otimes ... |n_{i-1}\rangle \otimes |n_i\rangle \otimes |n_{i+1}\rangle \otimes \ldots |n_N\rangle\,. \tag{S1.20}$$

##### S1.3 Derivation of a dynamical equation for population configurations

The dynamics of our individual-based model is stochastic. Hence, to fully characterize its dynamics, we must describe how the probability that the system is observed in a given configuration, $\eta$, changes with time. To this end, we first define a general lattice configuration at a time $t$, $|\psi(t)\rangle$, as the sum of all of the possible lattice configurations, i.e. the basis states, weighted by the probability of observing each of them

$$|\psi(t)\rangle = \sum_\eta P(\eta, t)\ |\eta\rangle\,, \tag{S1.21}$$

which changes in time according to

$$\frac{\partial}{\partial t} |\psi(t)\rangle = \sum_\eta \frac{\partial}{\partial t} P(\eta, t)\ |\eta\rangle\,. \tag{S1.22}$$

Therefore, the dynamics of the state $|\psi(t)\rangle$ is determined by the dynamics of the probabilities of finding the system in each of its possible states. These possible states are described by the basis states $|\eta\rangle$ and the probabilities of finding the system in each of them change according to the Master equation we derived in Section S1.1. Using some algebra and the commutation relationships for the bosonic operators in Eqs. (S1.16)-(S1.19), we can represent (S1.22) as a Schrodinger-like equation

$$\frac{\partial}{\partial t}\left|\psi(t)\right\rangle = -H(\{a^\dagger, a\})\left|\psi(t)\right\rangle \tag{S1.23}$$

with a quasi-hamiltonian given by $H(\{a^\dagger, a\}) = \sum_i \mathcal{H}_i(\{a^\dagger, a\})$ where

$$\mathcal{H}_i(\{a^\dagger, a\}) = b\left(\mathbb{I} - a_i^\dagger\right) a_i^\dagger a_i \,+\, d\left(a_i^\dagger - \mathbb{I}\right) a_i + \frac{\beta}{2R_f}\left(\mathbb{I} - a_i^\dagger\right) a_i^\dagger \sum_{j\,\in R_f(i)} \left\{a_j^\dagger a_j\right\} a_i \tag{S1.24}$$

$$+\,\frac{\gamma}{4R_c^2}\left(a_i^\dagger - \mathbb{I}\right) \sum_{j,k\,\in R_c(i)} \left\{a_j^\dagger a_k^\dagger a_k a_j\right\} a_i + \frac{h}{2}\sum_{\langle ij\rangle}\left(a_i^\dagger - a_j^\dagger\right)(a_i - a_j)\,. \tag{S1.25}$$

#### S1.4 Derivation of a path integral representation in terms of continuous field

To obtain a continuous equation for the dynamics of a population density field, we next need to derive a path integral representation of our model dynamics. First, we introduce the fields $\phi_i(t)$ and $\phi_i^*(t)$ in the Hamiltonian $\mathcal{H}(a^\dagger \to \phi(t)^* + 1,\ a \to \phi(t))$ and then take the continuum limit by letting the lattice spacing $\delta x \to 0$. To this end, we redefine the fields and parameters as

$$\frac{\phi_i(t)}{\delta x} \mapsto \phi(x,t), \quad \phi_i^*(t) \mapsto \tilde{\phi}(x,t), \quad \frac{h}{2} \mapsto \frac{D}{\delta x^2}, \tag{S1.26}$$

and the quasi-hamiltonian as $H\left(\phi^*(x,t), \phi(x,t)\right) = \int \mathcal{H}\left(\phi^*(x,t), \phi(x,t)\right) dx$. The new fields are associated with an action $S$, given by

$$S[\phi^*(x,t), \phi(x,t)] = \int dx \int_0^t d\tau \;\; \phi^* \partial_\tau \phi + \mathcal{H}[\phi^*(x,\tau), \phi(x,\tau)]. \tag{S1.27}$$

Next, we obtain the dynamic equation of the fields using the stationary-action principle or principle of least action, i.e. $\frac{\delta S}{\delta \phi} = \frac{\delta S}{\delta \tilde{\phi}} = 0$. By doing so, we obtain that the terms in the action that are linear in $\phi^*$ give rise to the equation

$$\frac{\partial \phi(x,t)}{\partial t} = \left[r + \frac{\beta}{2R_f}\int_{x-R_f}^{x+R_f} \phi(x',t)dx' - \frac{\gamma}{4R_c^2}\left(\int_{x-R_c}^{x+R_c} \phi(x',t)dx'\right)^2\right]\phi + D\,\nabla_x^2\phi. \tag{S1.28}$$

Finally, we take the expected value of the field, $\langle\phi\rangle$, which is equivalent to the mean density of particles $\rho$ in the continuum limit. Using the mean-field approximation $\langle\phi^2\rangle \approx \langle\phi\rangle^2 = \rho^2$, we obtain

$$\frac{\partial \rho(x,t)}{\partial t} = \left[r + \beta\,\tilde{\rho}_f(x,t) - \gamma\,\tilde{\rho}_c^2(x,t)\right]\rho(x,t) + D\,\nabla_x^2 \rho(x,t). \tag{S1.29}$$

where $\nabla_x^2$ is the Laplacian on the spatial coordinate $x$ and $\tilde{\rho}_\alpha(x,t)$ is the non-local density obtained by averaging the population density in the neighborhood of range $R_\alpha$ for $\alpha = \{f,\, c\}$

$$\tilde{\rho}_\alpha(x,t) = \int G\left(\left|x - x'\right|; R_\alpha\right)\rho\left(x',t\right)\mathrm{d}x'. \tag{S1.30}$$

$G(|x-x'|\,;R_\alpha)$ is the normalized interaction kernel for the facilitation and competition, $\alpha=\{f,\,c\}$,

$$G(\left|x-x'\right|;R_\alpha)=\begin{cases}\frac{1}{2R_\alpha} & \text{if } |x-x'|\le R_\alpha\\ 0 & \text{otherwise.}\end{cases}\tag{S1.31}$$

#### S2 Linear stability analysis of Eq. 2.5

In this section, we provide the detailed steps to perform a linear stability analysis of Eq. (S1.29), which gives the conditions to observe non-uniform spatial patterns of population density (Cross and Hohenberg, 1993). The linear stability analysis consists in introducing a small perturbation, $\epsilon\,\psi(x,t)$ with $\epsilon\ll 1$, to the uniform stable steady state of a partial differential equation and calculating the time evolution of the perturbation amplitude. If this amplitude decreases with time, the uniform state is also stable against spatial perturbations, and patterns do not form; if the perturbation amplitude increases, the uniform state is unstable against spatial perturbations, and spatial patterns form.

We consider a solution of the form $\rho(x,t)=\rho_+ + \epsilon\,\psi(x,t)$, where $\rho_+$ is the uniform stable state and $\psi$ is the perturbation of amplitude $\epsilon$. Inserting this solution in Eq. (S1.29) and retaining only linear terms in the perturbation, we get

$$\frac{\partial\psi(x,t)}{\partial t}=r\psi(x,t)+\beta\,\rho_+\Big[\psi(x,t)+\tilde{\psi}_f(x,t)\Big]-\gamma\,\rho_+^2\Big[\psi(x,t)+2\tilde{\psi}_c(x,t)\Big]+D\,\nabla^2\psi(x,t)\tag{S2.1}$$

where we have introduced the simplifying notation

$$\tilde{\psi}_\alpha(x,t)=\int G_\alpha(|x-a|)\,\psi(x,t)\,da.\tag{S2.2}$$

Equation (S2.1) is a linear integro-differential equation that we can solve using a Fourier transform:

$$\frac{\partial\hat{\psi}(k,t)}{\partial t}=\Big[r+\beta\,\rho_+\Big(1+\hat{G}_f(k)\Big)-\gamma\,\rho_+^2\Big(1+2\hat{G}_c(k)\Big)-Dk^2\Big]\hat{\psi}(k,t)\tag{S2.3}$$

where $\hat{\psi}(k,t)$ is the Fourier transform of the perturbation and $\hat{G}_\alpha(k)$ is the Fourier transform of the kernel

$$\hat{G}_\alpha(k)=\frac{\sin(R_\alpha k)}{R_\alpha\,k}.\tag{S2.4}$$

Finally, because Eq. (S2.3) is a linear differential equation, it has an exponential solution $\hat{\psi}(k,t)\propto\exp\left(\lambda(k)t\right)$ whose growth rate $\lambda(k)$ we can obtain from Eq. (S2.3)

$$\lambda(k)=\rho_+\Big[\beta\frac{\sin(R_f\,k)}{R_f\,k}-2\gamma\rho_+\frac{\sin(R_c\,k)}{R_c\,k}\Big]-Dk^2.\tag{S2.5}$$

We can also find the nondimensional counterpart of the growth-rate of the perturbation $\overline{\lambda}$, given by

$$\overline{\lambda}(\kappa)=u_+\Big[\frac{\sin(R\,\kappa)}{R\,\kappa}-2u_+\frac{\sin(\kappa)}{\kappa}\Big]-\overline{D}\kappa^2\tag{S2.6}$$

with the nondimensional wavenumber $\kappa = R_c k$.

#### S3 Numerical computation of the stationary states of Eq. 2.5

We compute the stationary states of Eq. 2.5 using its dimensionless counterpart in Eq. (S5.2). First, we integrate Eq. (S5.2) with $\overline{r} = 0$ until the population density reaches its stationary spatial pattern and compute the population abundance by integrating the density field over the system length. Next, we decrease $\overline{r}$ in a small amount $\Delta\overline{r}$ and integrate Eq. (S5.2) for a long time interval $\Delta\tau$ using the stationary pattern for $\overline{r} = 0$ as the initial condition. We repeat this process recursively until the $\overline{r}$ is such that the population density vanishes. This procedure gives us the stationary values of the population abundance that are stable, $\overline{u}_+$. To compute the unstable ones, we assume that $\overline{u}_-(\xi, \tau) = \alpha\,\overline{u}_+(\xi, \tau)$, where $\alpha$ is a dimensionless scaling parameter such that $0 < \alpha \leq 1$. This assumption implies that $\overline{u}_-$ has the same spatial structure as $\overline{u}_+$ and that both solutions only differ by a factor $\alpha$ that makes $\overline{u}_-(\xi, \tau) < \alpha\,\overline{u}_+(\xi, \tau)$. Under this assumption, to compute $\overline{u}_-$ we need to numerically find the $0 < \alpha \leq 1$ that satisfies $\partial_\tau \overline{u}_- = 0$. Because $\alpha = 1$ always satisfies this condition, we only take into account the lowest value of $\alpha$, which is $\alpha = 1$ only when $\overline{r} = \overline{r}_c$.

#### S4 Derivation of the group-level approximation in Eq. 3.2

We build this approximation to obtain estimates for the number of individuals within a single spatial aggregate of the stationary pattern using the three features of the spatial pattern discussed in the main text section 3.2:

1. All individuals within a group must interact with one another via competition and facilitation
2. Individuals of different groups must not compete with each other
3. If two groups interact with each other via facilitation, this positive interaction must reach all the individuals in both groups

First, we use the definition of $\tilde{\rho}$ in Eq. (S1.30). Given that the three assumptions above are fulfilled, $\tilde{\rho}_c$ is constant inside each aggregate. This is so because the integral that defines the averaged densities in Eq. (S1.30) is equal to the number of individuals within the group, $\mathcal{N}_i$. Therefore,

$$\tilde{\rho}_c(x_i, t) = \frac{\mathcal{N}_I(t)}{2R_c} \tag{S4.1}$$

where $x_i$ is the spatial coordinate of each lattice node, $i$, inside the aggregate "$I$". Following the same arguments, $\tilde{\rho}_f$ is also constant inside each group. However, since different groups can facilitate each other, the value of $\tilde{\rho}_f$ depends on the number of groups within the facilitation range $R_f$. Thus we can write

$$\tilde{\rho}_f(x_i, t) = \frac{\mathcal{N}_I(t)}{2R_f} + \sum_{\langle I,J\rangle} \frac{\mathcal{N}_J(t)}{2R_f} \tag{S4.2}$$

whereby $J$ runs over the neighbors of the focal group $I$ that are within the facilitation range. Next we use Eqs. (S4.1) and (S4.2) in Eq. (S1.29) to get:

$$\frac{\partial \rho(x_i,t)}{\partial t} = \left[ r + \beta \frac{\mathcal{N}_I(t)}{2R_f} + \beta \sum_{\langle I,J \rangle} \frac{\mathcal{N}_J(t)}{2R_f} - \gamma \frac{\mathcal{N}_I^2(t)}{4R_c^2} \right] \rho(x_i,t). \tag{S4.3}$$

where we have neglected the diffusion term because small diffusion is a necessary condition to have the three conditions above fulfilled. Finally, we integrate over the group length on both sides of Eq. (S4.3)

$$\frac{\partial \mathcal{N}_I(t)}{\partial t} = \left[ r + \frac{\beta}{2R_f} \left( \mathcal{N}_I(t) + \sum_{\langle I,J \rangle} \mathcal{N}_J(t) \right) - \gamma \frac{\mathcal{N}_I^2(t)}{4R_c^2} \right] \mathcal{N}_I(t). \tag{S4.4}$$

If we assume periodic boundary conditions, the spatial pattern of population density is periodic, and all aggregates have the same amount of neighbors and the same aggregate size. Hence, we can introduce a parameter that gives the number of groups within the facilitation range, $\eta$. Using this parameter, we can write an ordinary differential equation to describe the dynamics of aggregate size:

$$\frac{\partial \mathcal{N}(t)}{\partial t} = \left[ r + \beta(\eta+1) \frac{\mathcal{N}(t)}{2R_f} - \gamma \frac{\mathcal{N}^2(t)}{4R_c^2} \right] \mathcal{N}(t). \tag{S4.5}$$

Eq. (S4.5) is a cubic model with stationary solutions $\mathcal{N}_0 = 0$ and

$$\mathcal{N}_{\pm} = \frac{(\eta+1)\dfrac{\beta}{2R_f} \pm \sqrt{\left( (\eta+1)\dfrac{\beta}{2R_f} \right)^2 + \dfrac{r\gamma}{R_c^2}}}{\gamma/2R_c^2}, \tag{S4.6}$$

or, in its dimensionless form,

$$\mathcal{U}_{\pm} = \frac{(\eta+1)}{R} \pm \sqrt{\frac{(\eta+1)^2}{R^2} + 4r} \tag{S4.7}$$

where $\mathcal{U}_{\pm} = \gamma \mathcal{N}_{\pm}/\beta\, R_c$ given by

### S5 Model nondimensionalization

To facilitate the model analysis, we define scaled, dimensionless variables

$$u \equiv \frac{\gamma}{\beta}\rho \qquad \tau \equiv \frac{\beta^2}{\gamma} t \qquad \xi \equiv \frac{x}{R_c} \tag{S5.1}$$

which give dimensionless version of Eq. (S1.29):

$$\frac{\partial u(\xi,\tau)}{\partial \tau} = \left[ \overline{r} + \tilde{u}_f(\xi,\tau) - \tilde{u}_c^2(\xi,\tau) \right] u(\xi,\tau) + \overline{D}\, \nabla_\xi^2 u(\xi,\tau). \tag{S5.2}$$

where

$$\tilde{u}_f(\xi, \tau) = \int G\left(\left|\xi - \xi'\right|, R\right) u\left(\xi', \tau\right) \mathrm{d}\xi' \tag{S5.3}$$

and

$$\tilde{u}_c(\xi, \tau) = \int G\left(\left|\xi - \xi'\right|, 1\right) u\left(\xi', \tau\right) \mathrm{d}\xi' \tag{S5.4}$$

with scaled parameters

$$\overline{r} \equiv \frac{\gamma}{\beta^2} r \qquad \overline{D} \equiv \frac{\gamma}{(R_c\,\beta)^2} D \qquad R \equiv \frac{R_f}{R_c} \tag{S5.5}$$

Using this dimensionless model description, the total, non-dimensional population abundance is given by

$$\mathcal{A} = \frac{\gamma}{\beta\, R_c} A. \tag{S5.6}$$

### S6 Numerical methods

#### S6.1 Stochastic individual-based simulations

We simulate the stochastic individual-based model using the Gillespie algorithm, which is a widely used and efficient method to generate realizations of a stochastic process (Gillespie, 1976, 1977). The algorithm consists of two steps. First, we sample the time it takes for the next event to happen and, second, we sample which reaction takes place based on how each of them contributes to the total rate. The algorithm is based on the following steps:

1. At the beginning of the simulation, choose an initial condition for the number of individual in each lattice node.

2. Following Section S1.1, compute all the possible global transition rates $\Omega(\eta \to \eta')$ from the current configuration, $\eta$, to any other possible configuration $\eta'$. Define the total exit rate from the current configuration $\eta$,

$$\Omega_\eta^{\mathrm{OUT}} = \sum_{\eta'} \Omega(\eta \to \eta'). \tag{S6.1}$$

3. Sample the time to the next reaction, $\Delta t$, from an exponential distribution with mean equal to $1/\Omega_\eta^{\mathrm{OUT}}$.

4. Sample which of the possible transitions $\eta \to \eta'$ will take place. To do this sample, we define a probability of observing a transition to a specific configuration $\eta'$ as

$$P(\eta \to \eta') = \frac{\Omega(\eta \to \eta')}{\Omega_\eta^{\mathrm{OUT}}} \tag{S6.2}$$

5. Update the time and the configuration of the population to the new sampled values: $t \to t + \Delta t$ and $\eta \to \eta'$.

6. Repeat steps 2 to 5 until the desired simulation time is reached.

#### S6.2 Integration of the population-level approximation, Eq. 2.5

We perform the numerical integration of the population-level approximation PDE, Eq. 2.5 in the main text, using a second-order in time pseudospectral method detailed in Montagne et al. (1997). The method consists in determining the time evolution of a PDE in Fourier space through a "two-step" process after which we obtain the density field at a time $t+2\delta$ with an error $\mathcal{O}(\delta t^3)$. First, we Fourier transform the nonlinear PDE, Eq. 2.5 in the main text, and separate its linear and nonlinear terms

$$\frac{\partial\hat{\rho}(k,t)}{\partial t} = -\alpha(k)\hat{\rho}(k,t) + \Phi(k,t) \tag{S6.3}$$

where $\hat{\rho}(k,t)$ is the Fourier transform of the population density field and $\alpha(k) = Dk^2 - r$ is the coefficient associated with the linear part. $\Phi(k,t)$ is the Fourier transform of the nonlinear part of the original equation

$$\Phi(k,t) = \mathcal{F}\left[\tilde{\rho}_f(x,t)\rho(x,t) - \gamma\,\tilde{\rho}_c^2(x,t)\rho(x,t)\right] \tag{S6.4}$$

After setting the initial condition $\rho(x,0)$ and computing its Fourier transform $\hat{\rho}(k,0)$, the algorithm to integrate the equation between $t$ and $t+2\delta t$ is based on the following steps:

1. Compute $\Phi(x,t)$ in real space and transform it to Fourier space to obtain $\Phi(k,t)$.

2. Calculate the Fourier transform of the density field at time $t+\delta t$ (see Montagne et al. (1997) for a derivation of this expression) as,

$$\hat{\rho}(k,t+\delta t) = e^{-\alpha(k)\delta t}\hat{\rho}(k,t) + \frac{1-e^{-\alpha(k)\delta t}}{\alpha(k)}\Phi(k,t) \tag{S6.5}$$

3. Compute $\rho(x,t+\delta t)$ by Fourier transforming $\hat{\rho}(k,t+\delta t)$ and use it to calculate the nonlinear part of the original PDE in real space.

4. Compute $\Phi(k,t+\delta t)$ by Fourier transforming the result obtained in the previous step.

5. Calculate the updated field in the Fouier domain,

$$\hat{\rho}(k,t+2\delta t) = e^{-2\alpha(k)\delta t}\hat{\rho}(k,t) + \frac{1-e^{-2\alpha(k)\delta t}}{\alpha(k)}\Phi(k,t+\delta t) \tag{S6.6}$$

Thus, in each algorithm iteration, the field $\hat{\rho}(k,t)$ goes to $\hat{\rho}(k,t+2\delta t)$ and the process is repeated until the desired simulation time is reached. For all of the population-level model simulations, we use $dt = 0.05$, $dx = 0.008$.

#### S7 Supplementary Figure for the two-dimensional case

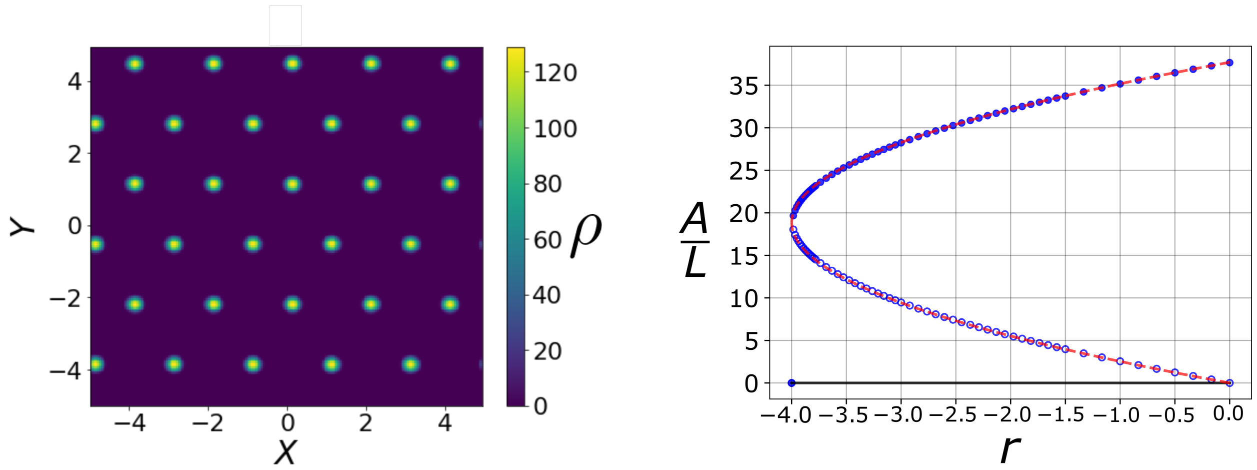

**Fig. S1:** Model results in two dimensions. a) Spatial pattern of population density for $r = -1$. (b) Total abundance divided by the system size in the presence of aggregates. Results are obtained using the deterministic equation for population density (blue circles) and the group-level approximation (dashed red line). Filled and empty symbols represent the stable and unstable states, respectively. The deterministic simulations run until $\tau = 3000$, $d\tau = 0.05$. $\beta = 1$, $\gamma = 1$, $R_f = 0.5$, $R_c = 1$ and $dx = 0.008$, $L = 10$ and $D = 10^{-3}$.

### References

Cross, M. C. and Hohenberg, P. C. (1993). Pattern formation outside of equilibrium. *Reviews of modern physics*, 65(3):851.

Doi, M. (1976). Stochastic theory of diffusion-controlled reaction. *Journal of Physics A: Mathematical and General*, 9(9):1479.

Gillespie, D. T. (1976). A general method for numerically simulating the stochastic time evolution of coupled chemical reactions. *Journal of computational physics*, 22(4):403–434.

Gillespie, D. T. (1977). Exact stochastic simulation of coupled chemical reactions. *The journal of physical chemistry*, 81(25):2340–2361.

Hernández-García, E. and López, C. (2004). Clustering, advection, and patterns in a model of population dynamics with neighborhood-dependent rates. *Physical Review E*, 70(1):016216.

Montagne, R., Hernández-García, E., Amengual, A., and San Miguel, M. (1997). Wound-up phase turbulence in the complex ginzburg-landau equation. *Physical Review E*, 56(1):151.

Peliti, L. (1985). Path integral approach to birth-death processes on a lattice. *Journal de Physique*, 46(9):1469–1483.

Täuber, U. C. (2007). *Field-Theory Approaches to Nonequilibrium Dynamics*, pages 295–348. Springer Berlin Heidelberg, Berlin, Heidelberg.
